# Cross-Tissue Comparison of Epigenetic Aging Clocks in Humans

**DOI:** 10.1101/2024.07.16.603774

**Authors:** Abner T. Apsley, Qiaofeng Ye, Avshalom Caspi, Christopher Chiaro, Laura Etzel, Waylon J. Hastings, Christine C. Heim, John Kozlosky, Jennie G. Noll, Hannah M. C. Schreier, Chad E. Shenk, Karen Sugden, Idan Shalev

**Affiliations:** Department of Biobehavioral Health, Penn State University, University Park, Pennsylvania, United States of America; Department of Molecular, Cellular, and Integrated Biosciences, The Pennsylvania State University, University Park, Pennsylvania, United States of America; Department of Psychology and Neuroscience, Duke University, Durham, North Carolina, United States of America; Social, Genetic and Developmental Psychiatry, Institute of Psychiatry, Psychology and Neuroscience, King’s College London, London, United Kingdom; PROMENTA, Department of Psychology, University of Oslo, Oslo, Norway; The Social Science Research Institute, Duke University, Durham, North Carolina, United States of America; Department of Psychiatry and Behavioral Science, Tulane University School of Medicine, New Orleans, Louisiana, United States of America; Charité–Universitätsmedizin Berlin, Corporate Member of Freie Universität Berlin, and Humboldt-Universitätzu Berlin, Berlin Institute of Health, Institute of Medical Psychology, Berlin, Germany; Department of Psychology, University of Rochester, Rochester, New York, United States of America; Department of Human Development and Family Studies, The Pennsylvania State University, University Park, Pennsylvania, United States of America; Department of Pediatrics, The Pennsylvania State University College of Medicine, Hershey, Pennsylvania, United States of America

## Abstract

Epigenetic clocks are a common group of tools used to measure biological aging – the progressive deterioration of cells, tissues and organs. Epigenetic clocks have been trained almost exclusively using blood-based tissues but there is growing interest in estimating epigenetic age using less-invasive oral-based tissues (i.e., buccal or saliva) in both research and commercial settings. However, differentiated cell types across body tissues exhibit unique DNA methylation landscapes and age-related alterations to the DNA methylome. Applying epigenetic clocks derived from blood-based tissues to estimate epigenetic age of oral-based tissues may introduce biases. We tested the within-person comparability of common epigenetic clocks across five tissue types: buccal epithelial, saliva, dry blood spots, buffy coat (i.e., leukocytes), and peripheral blood mononuclear cells. We tested 284 distinct tissue samples from 83 individuals aged 9-70 years. Overall, there were significant within-person differences in epigenetic clock estimates from oral-based versus blood-based tissues, with average differences of almost 30 years observed in some age clocks. In addition, most epigenetic clock estimates of blood-based tissues exhibited low correlation with estimates from oral-based tissues despite controlling for cellular proportions and other technical factors. Our findings indicate that application of blood-derived epigenetic clocks in oral-based tissues may not yield comparable estimates of epigenetic age, highlighting the need for careful consideration of tissue type when estimating epigenetic age.

## 1. Introduction

Measurements of biological age aim to capture the progressive deterioration of cells, tissues and organs (López-Otín et al. 2013; Schmauck-Medina et al. 2022). Over the past decade biological age measurements have provided new insights into the diverse trajectories of aging and related health risks that are often masked by reliance on chronological age. Among the varied methods proposed and tested for measuring biological age (Ferrucci et al. 2020), epigenetic age clocks – which are based on patterns of change in DNA methylation (DNAm) – stand out for their prevalence and utility (Salameh et al. 2020; Horvath & Raj 2018). These clocks have been applied widely across research fields, enhancing our understanding of aging in diverse contexts ranging from epidemiology to behavioral science. In particular, *epigenetic age acceleration*, when measured epigenetic age exceeds chronological age, has been associated with increased risk of multiple age-related diseases and early mortality (Fransquet et al. 2019).

Epigenetic clocks make use of the aging- and healthspan-related changes that take place in the DNA methylome by applying elastic-net regression methods over a wide range of DNAm measurements in order to predict a chosen outcome (Zou & Hastie 2005). Several generations of epigenetic clocks have been constructed via training on various outcomes such as chronological age in first-generation clocks (Horvath 2013; Hannum et al. 2013; McEwen et al. 2020; Horvath et al. 2018), phenotypic measurements of healthspan and time-to-death in second-generation clocks (Levine et al. 2018; Lu et al. 2022), and longitudinal physiological measurements of the pace of aging (Belsky et al. 2022). Parameters describing the association between DNAm measurements and a chosen outcome(s) in a training population are then applied to compute the epigenetic clock estimates of secondary samples.

Previous research has established wide intraindividual and interindividual variations in cellular composition and aging trajectories measured by DNAm (Hughes et al. 2015; Patrick et al. 2020; Santagata et al. 2014; Adalsteinsson et al. 2012; Theda et al. 2018). Each cell type has a unique DNAm signature (Kim & Costello 2017; Zilbauer et al. 2013; Hearn et al. 2019; van Dongen et al. 2018). Thus, estimates of epigenetic clocks can be skewed if cellular compositions vary across samples (Zhang et al. 2023) or if an epigenetic clock is computed in tissue types that differ from those in which the clocks were originally trained. For example, computing epigenetic clock estimates in saliva samples using clocks trained on blood-based tissues could result in different estimates of age acceleration due to differing DNAm signatures of blood-versus oral-based tissues. Furthermore, each tissue type can age at different rates within the same individual, thereby compounding the problem of measuring epigenetic clocks across varying tissue types and cellular compositions (Oh et al. 2023; Tian et al. 2023).

To overcome the problem of cell type heterogeneity, some epigenetic clocks have been constructed on multiple tissue types, such as the Horvath pan-tissue clock (Horvath 2013). However, most epigenetic clocks have been constructed using DNAm signatures of blood-based tissues such as whole blood (i.e., leukocytes) or peripheral blood mononuclear cells (PBMCs) (Hannum et al. 2013; Levine et al. 2018; Lu et al. 2022; Belsky et al. 2022). Given the challenges in blood collection for large population-based cohorts, there is a growing desire among researchers to use less invasive tissues, such as buccal or saliva (Raffington et al. 2021; Kim, Joyce, et al. 2023; Kim, Yaffe, et al. 2023; Milicic et al.; Chang & Lin 2023; Loh et al. 2023). These tissues do not require trained phlebotomists, can be collected in home settings, and there is less hesitancy among study participants to provide oral-based samples in comparison to having their blood drawn. In addition to academic research, commercial companies which offer epigenetic clock estimates for direct-to-consumer and health-care uses may prefer to use oral-based tissues to derive clock estimates that were developed in blood. Using oral-based tissues as the source of clock estimates could expand the applications of epigenetic clocks to more use-cases, both for consumers and for various health-care applications, such as identifying fast-aging patients to prescribe disease-preventive medications, or triaging slow-aging patients who are most likely to benefit from surgery (Safaee et al. 2023).

In research settings, oral-based tissue estimates of epigenetic clocks should (at minimum) be highly correlated with blood-based estimates (i.e., the rank order of estimates should be highly similar). High correlations between tissue-type estimates of epigenetic clocks enables the accurate testing of associations between estimates and phenotypes of interest. In contrast to research settings, commercial companies offering epigenetic clock estimates to customers need oral-based and blood-based tissue test results to agree absolutely. An oral-based clock estimate that incorrectly reports an individual to be epigenetically older or aging faster than expected is not a useful product.

Here, we tested the within-person comparability of epigenetic clocks across five tissue types collected in two cohort studies: buccal epithelial, saliva, dry blood spots (DBS), buffy coat (i.e., leukocytes), and PBMCs. Aiming to assess a wide range of clocks, we compared first-generation clocks: Horvath pan-tissue (Horvath 2013) and Hannum clocks (Hannum et al. 2013), second-generation clocks: PhenoAge (Levine et al. 2018) and GrimAge2 (Lu et al. 2022), and the DunedinPACE (Belsky et al. 2022). Additionally, we compared tissue estimates of the Skin and Blood clock (Horvath et al. 2018), due to having both blood- and oral-based tissues, and the PedBE clock (McEwen et al. 2020), which was constructed using buccal DNA in pediatric samples. We hypothesized that estimates of the Horvath pan-tissue clock would be similar across tissues (Horvath 2013), while clocks trained on a single type of tissue would differ across tissue types.

## 2. Materials & Methods

### 2.1 Participants and Design

Study participants were recruited from the Pennsylvania State University (PSU) community and surrounding areas, with some children recruited from other regions within Pennsylvania, as described in more detail below. Tissue samples were initially collected for cross-tissue comparisons of telomere length measurements (Wolf et al. 2024). Here, we used leftover tissue samples for cross-tissue comparisons of epigenetic clocks within-individuals. This study and protocols were approved by PSU’s Institutional Review Board.

#### 2.1.1 Adults

Adult participants were recruited via advertisements located on PSU’s University Park campus and in community bulletins in the surrounding areas. Approval from PSU’s Institutional Review Board was granted (protocol STUDY00008478), and all participants provided written informed consent. Inclusion criteria for the study included: (a) ages 18-75, (b) no significant medical illness or immune disease (e.g., cancer, diabetes, or autoimmune disease), (c) current non-smoker, and (d) not pregnant or currently breastfeeding. Individuals were excluded if they self-reported a recent infection, illness, and/or use of antibiotics. The maximum age was restricted to 75 years due to mortality selection (Cawthon et al. 2003). To balance across ages and sex, eligibility became more restricted as sampling progressed. Seventy-seven adults were recruited in total and the present investigation included the subset of 47 individuals who had at least two of the following tissues available: buccal, saliva, DBS and PBMC. Age (t(68)=1.31; p=0.19), sex (χ^2^(1)=0.16; p=0.69) and race (χ^2^(2)=0.53; p=0.77) were not significantly different between participants whose samples were included vs. excluded.

After obtaining informed consent, tissue samples and demographic information were collected from adult participants at PSU’s Clinical Research Center (CRC). Participants completed a set of questionnaires to collect demographic and health-related information. Trained phlebotomists performed antecubital venipuncture to collect 20 mL of whole blood in EDTA tubes. PBMCs were isolated from these whole blood samples through density-gradient centrifugation using Ficoll. Approximately 200 µL of whole blood were applied to a Whatman 903 protein saver card for the dried blood spot (DBS) samples. Participants were also asked to provide 4 mL of saliva across two Oragene tubes (OGR-500, DNA Genotek), which upon completion was mixed with Oragene stabilizing buffer and sealed. Eight buccal samples were collected using Isohelix SK1 swabs to firmly scrape the inside of the cheek per manufacturer’s directions. Collection order for all tissue types was uniform across participants. Participants were asked to refrain from eating or drinking anything other than water for one hour before arriving at the CRC. After collection, tissue samples were stored as follows: PBMCs were stored at −80°C in a solution buffer composed of phosphate buffered saline pH 7.2+EDTA (2mMol) + bovine serum albumin (0.5%) prior to DNA extraction. DBS were stored in sealed Ziploc bags with desiccant packets at room temperature. Buccal swabs were placed in sealed Ziploc bags and stored at −80°C. Saliva samples were aliquoted into 4 cryovials and stored at −80°C.

#### 2.1.2 Children

Child participants were members of the Child Health Study (CHS), a large cohort study designed to provide prospective, longitudinal data on the health and development of children with and without a history of child maltreatment investigations (for more details about the CHS see Schreier et al. 2021). Approval from PSU’s Institutional Review Board was granted (protocol STUDY00006550), and informed assent (child) and consent (caregiver) was obtained for all participants. The CHS is actively following a cohort of 700 children and the present investigation included a random subset of 36 children who had at least two of the following tissues available: buccal, saliva, DBS and buffy coat. Age was significantly higher in the included samples (t(42)=3.37; p=0.002), but both sex (χ^2^(3)=2.93; p=0.40) and race (χ^2^(5)=4.21; p=0.52) were not significantly different between included and excluded samples.

Caregivers accompanied children to PSU’s University Park campus. Tissue samples were collected from the child participants and their caregivers provided information on child health and demographics. Subsequently, trained phlebotomists collected 20 mL of whole blood in EDTA tubes via antecubital venipuncture from youth. Buffy coat was isolated using centrifugation to separate plasma followed by treatment with 0.5x red blood cell lysis buffer (Invitrogen). Using identical procedures to those described in adults, approximately 200 µL of whole blood were used to collect a DBS sample on a Whatman 903 protein saver card; two mL of saliva (Oragene OGR-500, DNA Genotek) and two buccal swabs (Isohelix SK1) were also taken per individual. DBS, saliva, and buccal swabs were stored in the same conditions as adult samples, and buffy coat was stored at −80°C in a solution buffer composed of phosphate buffered saline pH 7.2+EDTA (2mMol) + bovine serum albumin (0.5%).

#### 2.1.3 Demographic Measures

Chronological age, biological sex, and race/ethnicity were included as covariates. All demographic variables were measured via self-report.

### 2.2 DNA Extraction

To minimize the impact of DNA extraction procedures, DNA was extracted from all tissue samples using the Gentra Puregene DNA Extraction Kit according to factory guidelines (Qiagen). This kit has been used to extract DNA from whole blood, PBMCs, saliva, buccal cells, and DBS (Koontz et al. 2015). Extracted DNA was stored at −80°C in Qiagen DNA Hydration Solution.

### 2.3 DNA Methylation Measurements

DNA (N=83 individuals, N=296 total samples) was delivered to PSU’s Genomics Core Facility for bisulfite conversion, processing, and methylation measurements. All bisulfite conversions, DNA processing, and array hybridization steps were performed by the same technician to decrease technical variability. The Infinium MethylationEPIC v2.0 BeadChip Kit was used to measure DNA methylation in over 935,000 CpG sites across the genome and four array plates were used in total (N_1_=96; N_2_=96; N_3_=96; N_4_=8). IDAT files were read into R statistical software using the *read.metharray.exp* function in the *minfi* package (Aryee et al. 2014). A total of 12 tissue samples (10 saliva and 2 buffy coat) across 11 individuals with probe detection p-values greater than 0.05 for more than 5% of probes or with bisulfite conversion rates of less than 80% were excluded from downstream analyses leaving 284 tissue samples from 83 individuals. CpG probes with a bead count of less than 3 in 5% or more samples or with an average detection p-value of > 0.05 across all samples were excluded (# of probes excluded = 8,419). Prior to calculation of epigenetic clocks, all samples were normalized using the beta-mixture quantile normalization (BMIQ) method (Teschendorff et al. 2013).

### 2.4 CpG Probe Imputation

All clocks included in our downstream analyses contained CpG probes that are not present on the Infinium MethylationEPIC v2.0 BeadChip Array (Kaur et al. 2023). The number of probes present for each clock is shown in supplementary **Table S1**. Missing probes were imputed using custom “golden standard” datasets constructed for each tissue type and age (1-18% imputation per clock). For further information on imputation methods, see **Supplementary Materials** and **Table S2**.

### 2.5 Epigenetic Clock Estimates

Horvath pan-tissue, Hannum, PhenoAge, GrimAge2, DunedinPACE, Skin and Blood, and PedBE estimates were calculated manually using modified code from both the *methylCIPHER* (Thrush et al. 2022) and the *DunedinPACE* packages (Belsky et al. 2022) in R. To perform sensitivity analyses, principal component (PC) clock measurements of the Horvath pan-tissue, Hannum, PhenoAge, GrimAge and Skin and Blood clocks were computed manually using source code provided by the authors of these clocks (Higgins-Chen et al. 2022). Due to the robustness of PC clocks to missing probes, no imputation was performed on missing CpG sites when computing these clocks. Epigenetic age acceleration measurements were computed for each clock (excluding DunedinPACE) as the difference between epigenetic age and chronological age in years.

### 2.6 Estimation of Cellular Composition of Each Tissue

Cellular compositions for buccal and saliva tissues were estimated using methods described by Houseman et al. (Houseman et al. 2016). Briefly, a reference-free cellular decomposition method was applied separately to all buccal and saliva samples. In order to determine the number of cell subtypes that should be used for each tissue, the top 10,000 most variable probes across all samples were extracted separately from buccal and saliva tissue. Next, for estimates of 1-15 cell subtypes for buccal and saliva tissues, a deviance statistic was calculated for 1000 bootstrapped samples. The number of cell subtypes was chosen to minimize the quantile-trimmed mean deviance statistics for both buccal and saliva, with buccal having 3 cellular subtypes and saliva having 5 cellular subtypes (see **Figure S1**).

DBS, buffy coat, and PBMC cellular subtype compositions were estimated using DNAm estimates of immune cell proportions (Houseman et al. 2012). DBS and buffy coat cellular estimates included CD4T, CD8T, natural killer, B-cell, monocyte and granulocyte estimates. PBMC cellular estimates included CD4T, CD8T, natural killer, B-cell and monocyte estimates.

### 2.7 Statistical Analyses

Statistical analyses were performed using R Studio v.2023.06.2 (R 4.3.1). Epigenetic clock estimates were stratified by tissue (buccal, saliva, DBS, buffy coat and PBMC) and displayed with violin plots using *ggplot2*. Within-person Pearson bivariate correlations for all clock acceleration values were calculated for all tissue pairs and displayed in correlation heatmaps using *corrplot*. Within-person differences between clock estimates were calculated using a paired t-test for each tissue pair. Intraclass correlation coefficients (ICCs) were computed across all tissues using the *ICC* function in the *psych* package.

We conducted two sensitivity analyses. The first consisted of computing PC epigenetic clock estimates for the Horvath pan-tissue, Hannum, PhenoAge, GrimAge and Skin and Blood clocks, and analyzing within-person differences between PC estimates for each set of tissues using paired t-tests. Secondly, we computed within-person partial Pearson correlations for all clock acceleration values across tissues, controlling for cellular compositions of tissues (see *Section 2.6*) and DNAm batch to determine whether cellular composition of tissues and DNAm batch altered the correlations between epigenetic age accelerations. DNAm batch residualization was performed using the first 30 principal components (greater than 70% of the variance) of control-probe beta values on the Infinium MethylationEPIC v2.0 BeadChip Array (Lehne et al. 2015). Because all comparisons were made within-person, sex and race/ethnicity were not included as covariates.

## 3. Results

### 3.1 Descriptive Statistics

Our analytical sample consisted of 83 individuals across two studies with a total of 284 tissue samples (N_Buccal_=81; N_Saliva_=59; N_DBS_=64; N_BuffyCoat_=35; N_PBMC_=45). Seventy-five individuals had within-person measurements for at least three tissue types with over half of individuals having four tissue types (N_2-Tissue_=8; N_3-Tissue_=32; N_4-Tissue_=43). Two tissue types differed between cohorts (i.e., buffy coat was collected only in children and PBMCs were collected only in adults). Demographic statistics for the sample, stratified by cohort, are shown in

**Table 1:**
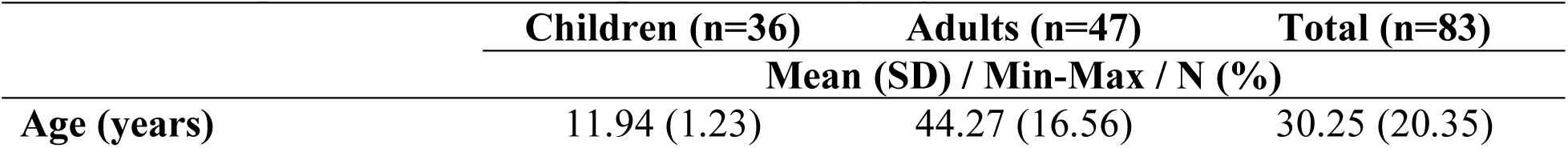
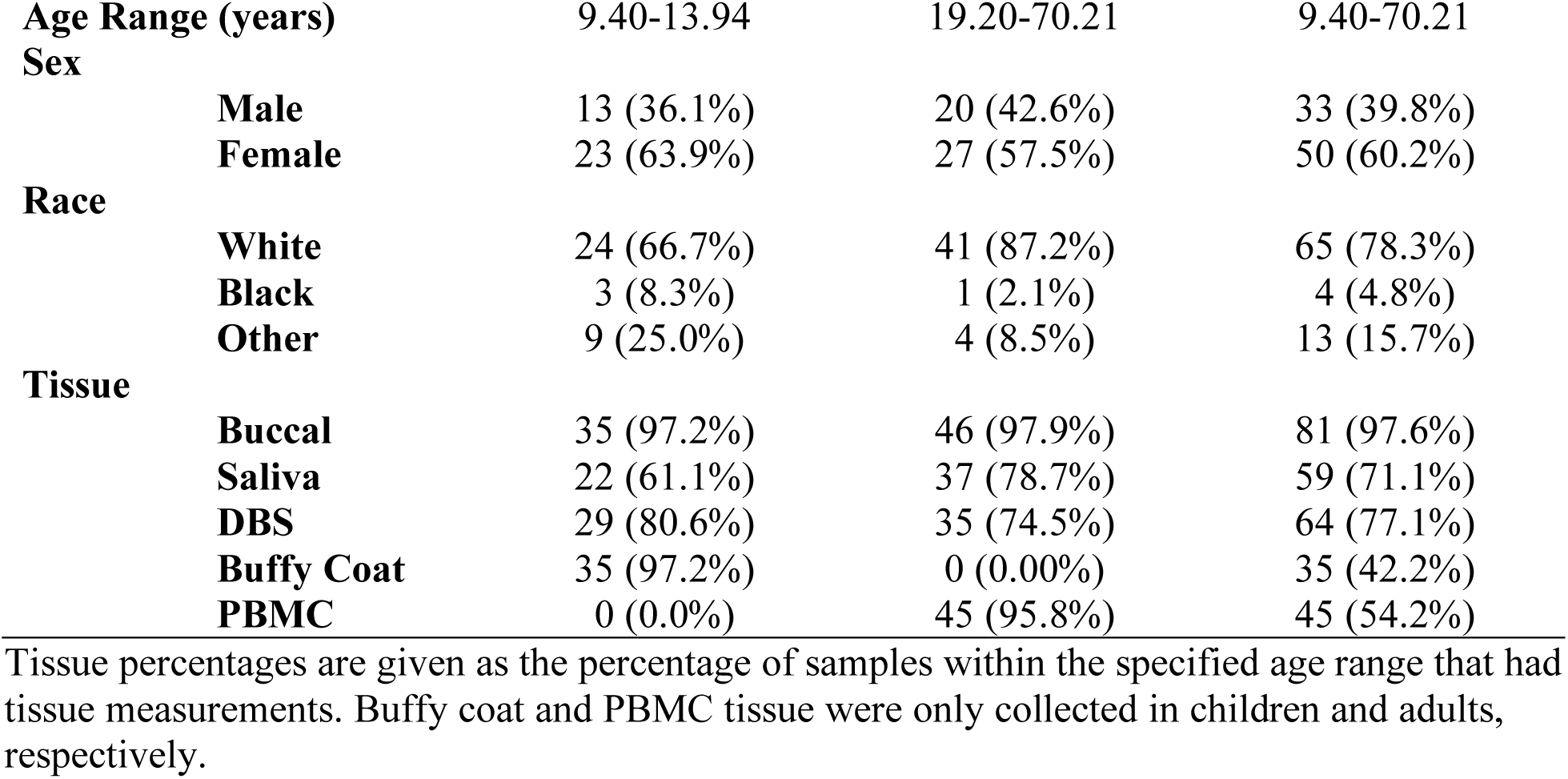
Descriptive Statistics of Sample Stratified by Age.

### 3.2 Comparability of First-Generation Epigenetic Clocks Across Tissues

First-generation epigenetic clock estimates varied across tissues. Moreover, the directionality of differences between tissues was not maintained between the Horvath pan-tissue and Hannum epigenetic age clocks.

Mean Horvath pan-tissue epigenetic age estimates were highest for saliva samples (43.44 ± 21.59) and lowest for buccal (32.08 ± 20.67) (**Figure 1a**). As expected, mean values of Horvath pan-tissue ages for buffy coat (23.05 ± 4.48) and PBMCs (53.74 ± 16.87) were different due to the restricted age ranges of samples with each tissue type. For Horvath pan-tissue age acceleration measurements, positive mean values were observed in all tissues (Buccal=1.60 ± 4.39; Saliva=11.15 ± 7.38; DBS=13.12 ± 4.07; Buffy coat=11.18 ± 3.87; PBMCs=9.99 ± 4.95), indicating higher epigenetic age estimates than chronological age (**Figure 1b**). Within-person correlations of Horvath pan-tissue age acceleration measurements were strongest for DBS and buffy coat (*r*=0.74) and weakest for saliva and PBMC (*r*=0.00) (**Figure 1c**).

**Figure 1.**
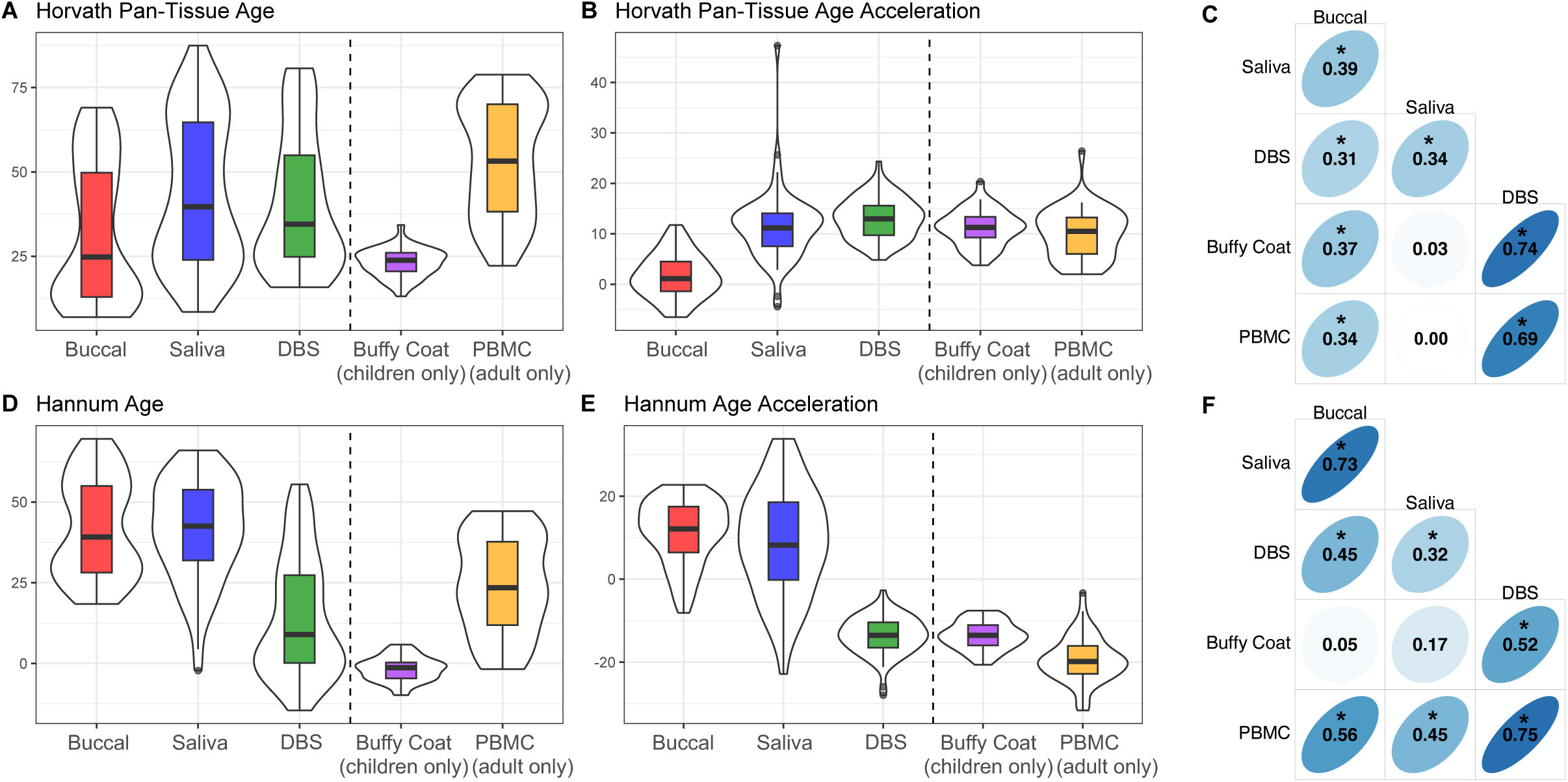
First Generation Epigenetic Clock Distributions and Correlations. (A) Horvath pan-tissue epigenetic age estimates, (B) Horvath pan-tissue epigenetic age acceleration estimates, (C) within-person correlations of Horvath pan-tissue epigenetic age acceleration across tissues, (D) Hannum epigenetic age estimates, (E) Hannum epigenetic age acceleration estimates, and (F) within-person correlations of Hannum epigenetic age acceleration across tissues. Vertical dashed lines in (A), (B), (D) and (E) separate tissues measured in both adults and children (buccal, saliva and DBS) and tissues measured in either adults (PBMCs) or children (buffy coat). Thick black horizontal bars on violin plots indicate the tissue-stratified median clock value and colored boxes indicate interquartile ranges. * indicates correlations with p<0.05.

Hannum age estimates were highest for buccal (41.76 ± 15.18) and saliva samples (41.16 ± 15.46), whereas DBS exhibited a lower mean value (14.46 ± 18.36) (**Figure 1d**). As expected, mean values of Hannum age for adult PBMCs (24.27 ± 15.12) were higher than for child buffy coat samples (−1.73 ± 3.88), the latter of which were registered as having negative epigenetic age in most samples. For Hannum age acceleration estimates, positive values were observed in buccal (11.27 ± 7.77) and saliva (8.87 ± 12.88) tissues, whereas negative values were observed in DBS (−13.56 ± 4.52), buffy coat (−13.60 ± 3.44) and PBMCs (−19.47 ± 5.57), indicating higher epigenetic age estimates than chronological age in buccal and saliva and lower epigenetic age estimates than chronological age in DBS, buffy coat, and PBMCs (**Figure 1e**). Within-person correlations of Hannum age acceleration measurements were strongest for DBS and PBMCs (*r*=0.75) and weakest for buccal and buffy coat (*r*=0.05) (**Figure 1f**).

Mean, median and standard deviations of both Horvath pan-tissue and Hannum estimates are detailed in supplementary **Table S3**. Results of first-generation epigenetic clocks across all tissue types stratified by age are provided in supplementary **Figures S2** and **S3**. ICCs across all tissues for both Horvath pan-tissue and Hannum estimates are detailed in supplementary **Table S4**.

### 3.3 Comparability of Second-Generation Epigenetic Clocks Across Tissues

Second-generation epigenetic clock estimates also varied across tissues, however, the directionality of differences was more consistent than was observed for first-generation clocks.

PhenoAge estimates for buccal (43.94 ± 18.85) and saliva (50.31 ± 22.98) had the highest mean values, whereas DBS had the lowest (21.70 ± 22.17) (**Figure 2a**). As expected, mean values of buffy coat (3.42 ± 5.83) and PBMCs (29.03 ± 18.91) PhenoAge measurements were different due to the restricted age ranges of participants who provided samples with each tissue type. We observed negative mean values for PhenoAge acceleration measurements in DBS (−6.31 ± 5.98), buffy coat (−8.44 ± 5.38) and PBMCs (−14.72 ± 6.68), and positive values in both buccal (13.45 ± 6.19) and saliva (18.02 ± 12.27) (**Figure 2b**). Within-person correlations of PhenoAge acceleration measurements were strongest for DBS and buffy coat (*r*=0.75) and weakest for buccal and DBS (*r*=0.05) (**Figure 2c**).

**Figure 2.**
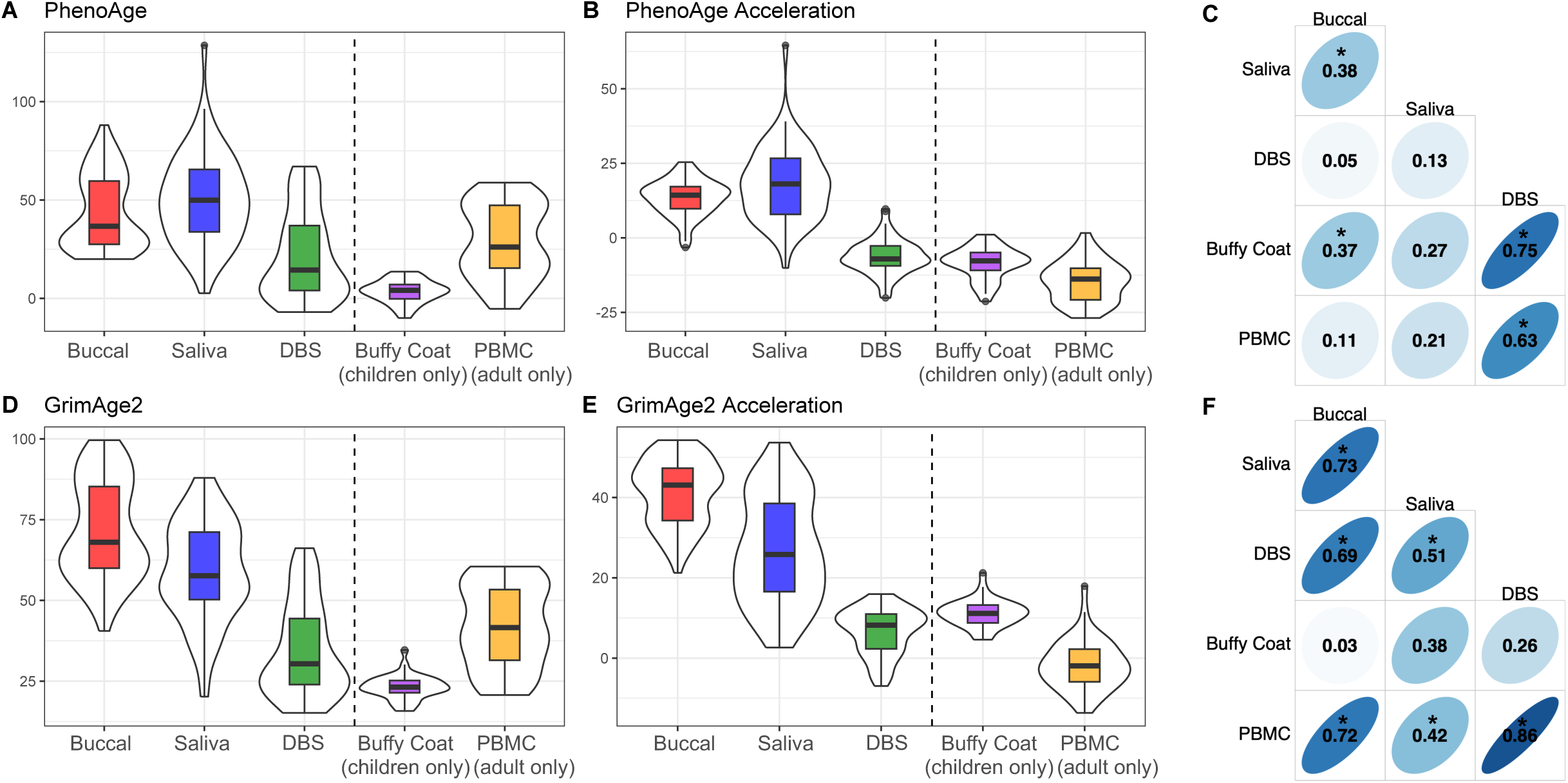
Second Generation Epigenetic Clock Distributions and Correlations. (A) PhenoAge epigenetic age estimates, (B) PhenoAge epigenetic age acceleration estimates, (C) within-person correlations of PhenoAge epigenetic age acceleration across tissues, (D) GrimAge2 epigenetic age estimates, (E) GrimAge2 epigenetic age acceleration estimates, and (F) within-person correlations of GrimAge2 epigenetic age acceleration across tissues. Vertical dashed lines in (A), (B), (D) and (E) separate tissues measured in both adults and children (buccal, saliva and DBS) and tissues measured in either adults (PBMCs) or children (buffy coat). Thick black horizontal bars on violin plots indicate the tissue-stratified median clock value and colored boxes indicate interquartile ranges. * indicates correlations with p<0.05.

GrimAge2 estimates were also highest for buccal and saliva (Buccal=71.68 ± 15.36; Saliva=59.17 ± 15.27) and lower for DBS (34.69 ± 14.50) (**Figure 2d**). GrimAge2 estimates of child buffy coat samples were higher than expected (23.14 ± 3.89) but were still well below estimates of adult PBMCs (42.01 ± 12.73), in line with the restricted age ranges of participants who provided samples with each tissue type. GrimAge2 acceleration estimates showed higher mean values for buccal (41.19 ± 8.10) and saliva (26.88 ± 13.36) tissues, and lower mean value for DBS (6.67 ± 5.73), and buffy coat (11.27 ± 3.43). Adult PBMCs were the only tissue to exhibit negative GrimAge2 acceleration estimates (−1.74 ± 6.15) (**Figure 2e**). Within-person correlations of GrimAge2 acceleration measurements were strongest for DBS and PBMCs (*r*=0.86) and weakest for buccal and buffy coat (*r*=0.03) (**Figure 2f**).

Mean, median and standard deviations of both PhenoAge and GrimAge2 estimates are detailed in supplementary **Table S5**. For second-generation epigenetic clocks across all tissue types, stratified by age, see supplementary **Figures S4** and **S5**. ICCs across all tissues for both PhenoAge and GrimAge2 estimates are detailed in supplementary **Table S6**.

### 3.4 Comparability of DunedinPACE Epigenetic Clock Across Tissues

DunedinPACE estimates varied, with buccal and saliva exhibiting the highest rate of epigenetic aging (Buccal=1.61 ± 0.08; Saliva=1.50 ± 0.25) and DBS (0.96 ± 0.09), buffy coat (0.96 ± 0.09) and PBMCs (0.89 ± 0.10) all exhibiting lower rates (**Figure 3a**). Within-person correlations of DunedinPACE measurements (**Figure 3b**) were strongest for DBS and buffy coat (*r*=0.72) and weakest for saliva and PBMC (*r*=0.15).

**Figure 3.**
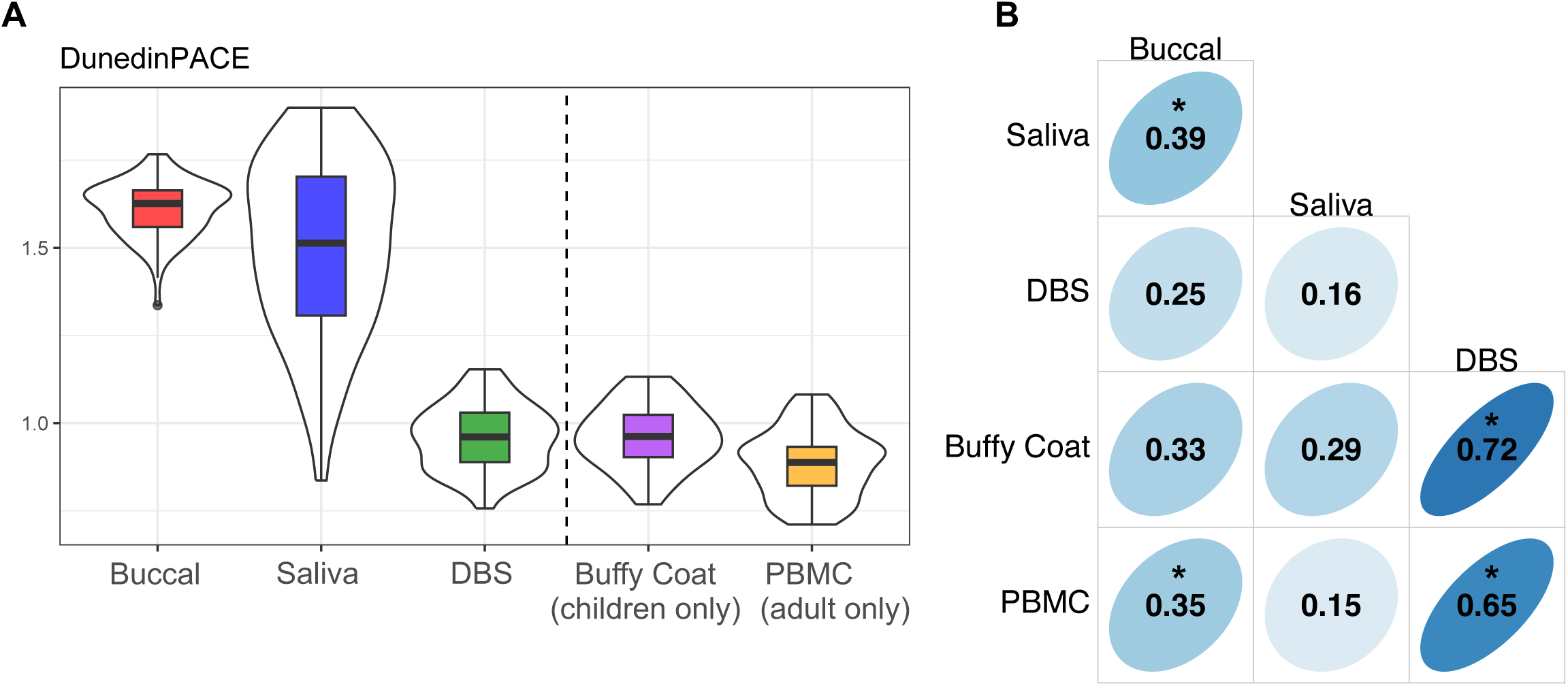
DunedinPACE Epigenetic Clock Distributions and Correlations. (A) DunedinPACE estimates, (B) within-person correlations of DunedinPACE across tissues. Vertical dashed lines in (A) separate tissues measured in both adults and children (buccal, saliva and DBS) and tissues measured in either adults (PBMCs) or children (buffy coat). Thick black horizontal bars on violin plots indicate the tissue-stratified median clock value and colored boxes indicate interquartile ranges. * indicates correlations with p<0.05.

Mean, median and standard deviations of DunedinPACE estimates are detailed in supplementary **Table S7**. For DunedinPACE estimates across all tissue types, stratified by age, see supplementary **Figures S6** and **S7**. ICCs across all tissues for DunedinPACE estimates are detailed in supplementary **Table S8**.

### 3.5 Comparability of Skin and Blood and PedBE Epigenetic Clocks Across Tissues

Skin and Blood clock age estimations were less variable across tissues. Even so, saliva (32.09 ± 20.80) and buccal (28.65 ± 20.32) estimates tended to be higher than estimates from DBS (25.39 ± 19.45) (**Figure 4a**). As expected, mean values of buffy coat (8.95 ± 1.44) and PBMCs (40.48 ± 16.25) Skin and Blood age were different due to the restricted age ranges of participants who provided samples with each tissue type. For Skin and Blood age acceleration measurements, all tissues exhibited negative values (Buccal=−1.84 ± 3.41; Saliva=−0.21 ± 0.62; DBS=−2.63 ± 2.36; Buffy coat=−2.92 ± 1.25; PBMCs=−3.26 ± 3.53) (**Figure 4b**). Within-person correlations of Skin and Blood acceleration measurements were strongest for DBS and buffy coat (*r*=0.68) and weakest for saliva and buffy coat (*r*=−0.11) (**Figure 4c**).

**Figure 4.**
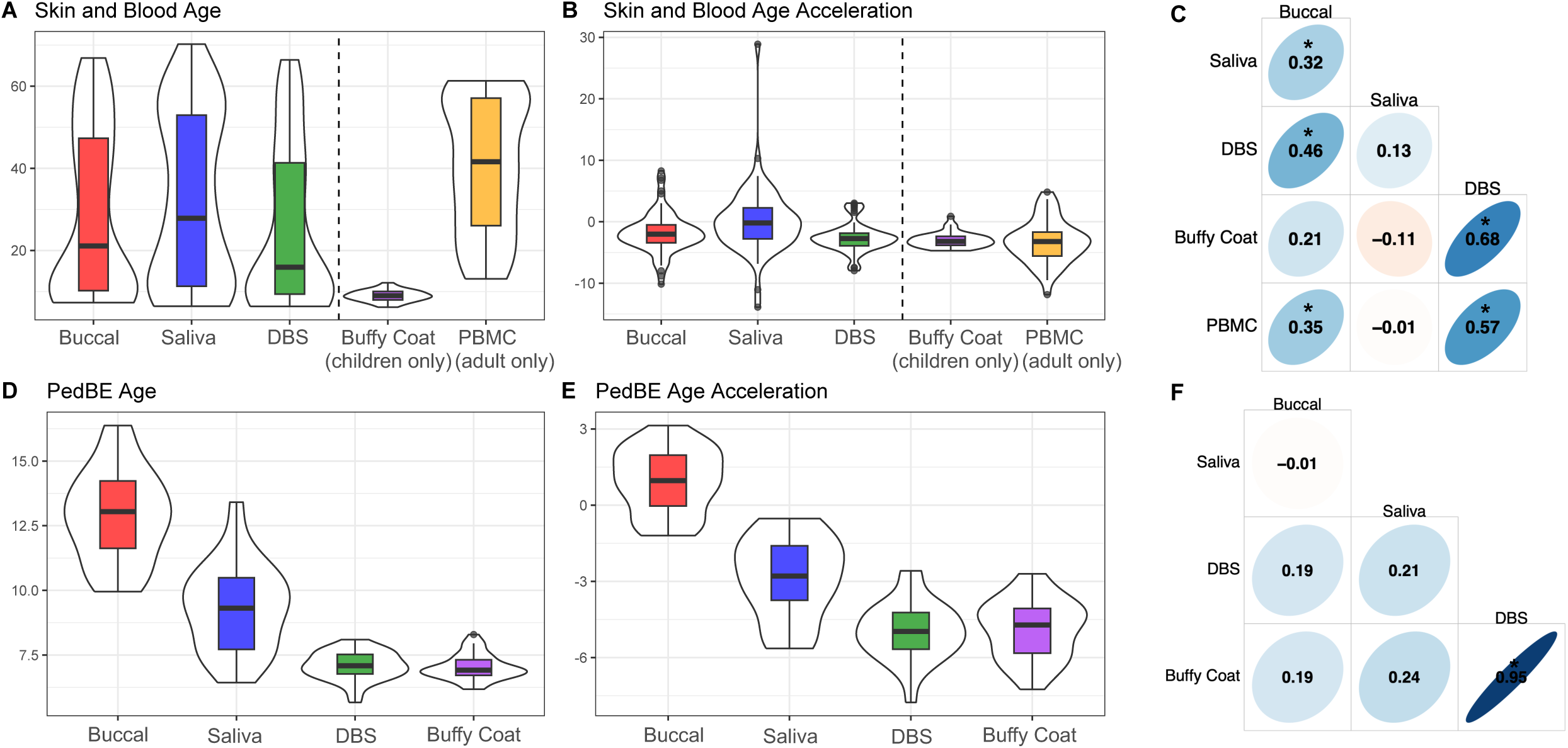
Skin and Blood and PedBE Clock Distributions and Correlations. (A) Skin and Blood epigenetic age estimates, (B) Skin and Blood epigenetic age acceleration estimates, (C) within-person correlations of Skin and Blood epigenetic age acceleration across tissues, (D) PedBE epigenetic age estimates, (E) PedBE epigenetic age acceleration estimates, and (F) within-person correlations of PedBE epigenetic age acceleration across tissues. Vertical dashed lines in A and B separate tissues measured in both adults and children (buccal, saliva and DBS) and tissues measured in either adults (PBMCs) or children (buffy coat). Thick black horizontal bars on violin plots indicate the tissue-stratified median clock value and colored boxes indicate interquartile ranges. * indicates correlations with p<0.05.

PedBE age estimates were derived within child samples only. Buccal and saliva tissues exhibited higher mean values (Buccal=12.89 ± 1.77; Saliva=9.32 ± 1.69), whereas estimates from DBS and buffy coat tissues were lower and in close alignment with one another (DBS=7.087± 0.55; Buffy Coat=7.03 ± 0.49) (**Figure 4d**). PedBE acceleration estimates were positive for buccal (0.97 ± 1.25) but were negative for all other tissues (Saliva=−2.86 ± 1.63; DBS=−4.97 ± 1.12; Buffy Coat=−4.84 ± 1.14) (**Figure 4e**). Within-person correlations of PedBE acceleration measurements were strongest for DBS and buffy coat (*r*=0.95) and weakest for buccal and saliva (*r*=−0.01) (**Figure 4f**).

Mean, median and standard deviations of both Skin and Blood and PedBE estimates are detailed in supplementary **Table S9**. Skin and Blood and clock measurements across all tissue types stratified by study cohort are shown in supplementary **Figures S8** and **S9**. ICCs across all tissues for both Skin and Blood and PedBE estimates are detailed in supplementary **Table S10**.

### 3.6 Pairwise Within-Person Comparisons of Epigenetic Clock Estimates

Within-person differences in epigenetic clock estimates were most pronounced between oral and blood-based tissues. All within-person clock estimates, other than Horvath pan-tissue, were significantly higher in buccal and saliva than in blood-based tissues (i.e., DBS, PBMCs, buffy coat; differences ranging from 0.84 for Skin and Blood to 40.25 for GrimAge2; see **Table 2**). In contrast, Horvath pan-tissue clock estimates for buccal tissue were lower than estimates for all blood tissues (differences ranging from 8.12-11.09 years), and differences between saliva and blood tissues were not significant (differences ranging from −1.88 to 2.22 years).

**Table 2.** Pairwise Within-Person Differences in Standard Epigenetic Clocks Across Tissue Types.

|  | Buccal vs Saliva | Buccal vs DBS | Buccal vs Buffy Coat | Buccal vs PBMC | Saliva vs DBS | Saliva vs Buffy Coat | Saliva vs PBMC | DBS vs Buffy Coat | DBS vs PBMC |
| --- | --- | --- | --- | --- | --- | --- | --- | --- | --- |
| | $\beta \pm \text{SE}$ | | | | | | | | |
| **Horvath** | -9.45 ± 0.96*** | -11.09 ± 0.65*** | -10.12 ± 0.71*** | -8.12 ± 0.90*** | -1.88 ± 1.13 | -1.69 ± 1.44 | 2.22 ± 1.69 | -0.27 ± 0.56 | 4.33 ± 0.66*** |
| **Hannum** | 1.29 ± 1.19 | 26.18 ± 0.80*** | 29.29 ± 1.04*** | 28.26 ± 1.06*** | 23.28 ± 1.71*** | 29.28 ± 2.96*** | 24.16 ± 1.71*** | 0.02 ± 0.80 | 5.46 ± 0.66*** |
| **PhenoAge** | -5.33 ± 1.51*** | 20.96 ± 1.02 *** | 23.98 ± 1.01*** | 27.10 ± 1.43*** | 24.54 ± 2.05*** | 24.93 ± 2.70*** | 32.75 ± 2.25*** | -1.33 ± 0.78 | 11.23 ± 0.98*** |
| **GrimAge2** | 13.47 ± 1.23*** | 35.78 ± 0.70*** | 34.82 ± 1.28*** | 40.25 ± 0.82*** | 20.72 ± 1.73*** | 25.70 ± 2.62*** | 22.42 ± 1.63*** | -1.11 ± 0.84 | 5.16 ± 0.58*** |
| **DunedinPACE** | 0.10 ± 0.03** | 0.65 ± 0.01*** | 0.62 ± 0.02*** | 0.75 ± 0.01*** | 0.53 ± 0.04*** | 0.61 ± 0.06*** | 0.58 ± 0.04*** | -0.02 ± 0.01 | 0.10 ± 0.01*** |
| **Skin and Blood** | -1.91 ± 0.83* | 1.33 ± 0.39*** | 0.84 ± 0.31** | 2.01 ± 0.70** | 2.85 ± 1.05** | 1.32 ± 0.69 | 3.98 ± 1.46*** | -0.29 ± 0.19 | 0.86 ± 0.55 |
| **PedBE** | 3.89 ± 0.49*** | 5.90 ± 0.30*** | 5.90 ± 0.28*** | -- | 1.99 ± 0.42*** | 2.20 ± 0.40*** | -- | 0.04 ± 0.07 | -- |

P < 0.05 \*, P < 0.01 \*\*, P < 0.001 \*\*\*

Although Hannum estimates were similar between buccal and saliva, estimates of all other epigenetic clocks exhibited varying degrees of concordance, with buccal having higher estimates than saliva in GrimAge2 (13.47 ± 1.23) and PedBE (3.89 ± 0.49) clocks, and lower estimates than saliva in Horvath pan-tissue (9.45 ± 0.96), PhenoAge (5.33 ± 1.51), and Skin and Blood (1.91 ± 0.83) clocks. Buccal tissues also displayed faster rates of aging than saliva using the DunedinPACE epigenetic clock (0.10 ± 0.03). No significant differences between DBS and buffy coat were observed for any clock. In contrast, in all clocks tested, DBS had higher clock estimates when compared to PBMCs (differences ranging from 0.86-11.23), although these differences did not reach statistical significance for the Skin and Blood clock. No comparisons were made between buffy coat and PBMCs because these samples were obtained from two different studies. Age-stratified tissue comparisons are reported in supplementary **Tables S11** and **S12.**

### 3.7 Sensitivity Analyses – PC Clocks and Controlling for Cellular Composition and DNA Methylation Batch

Although values for within-person, between-tissue comparisons are different for standard and PC clocks (compare **Table 2** and supplementary **Table S13**), the direction of effects and significance levels of PC clocks for most tissues and clocks remained comparable to the effects observed when using standard clocks. Comparisons between standard and PC clock estimates within-person and within-tissue are shown in supplementary **Table S14**. Violin plots of PC clock estimates (full sample and age-stratified) are shown in supplementary **Figures S10-S18**.

To estimate the effect of cellular composition and DNAm batch, we residualized epigenetic age acceleration values by both factors to determine whether cellular composition or batch influenced epigenetic age acceleration correlations. Overall, when controlling for cell composition and DNAm batch, a majority of correlations decreased. Notable improvements were observed in Skin and Blood clock saliva-buffy coat correlations (*r*=−0.11 to *r*=0.27) and Skin and Blood saliva-PBMC correlations (*r*=−0.01 to 0.17) (**Figure 5**).

**Figure 5.**
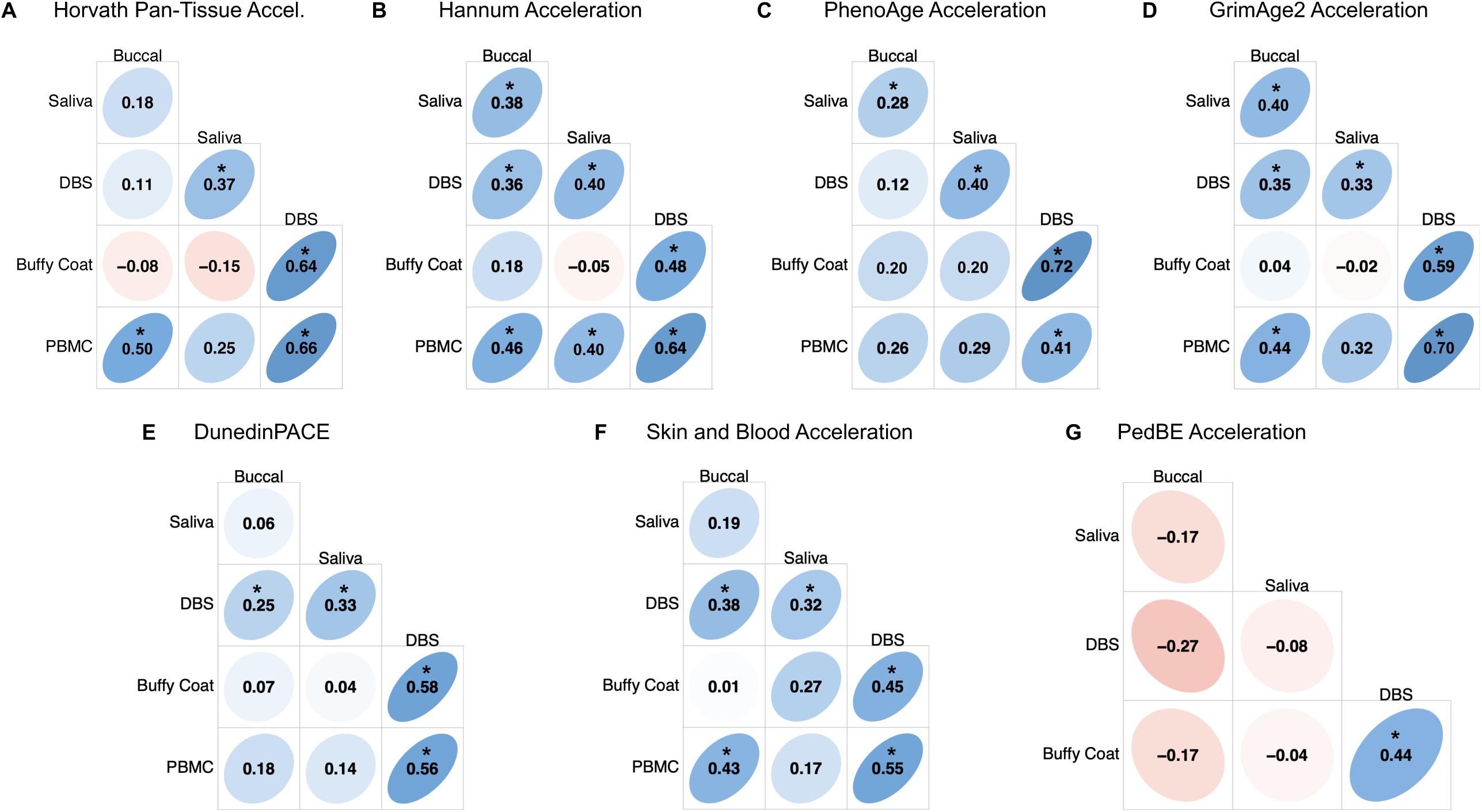
Within-Person Epigenetic Age Acceleration Correlations Across Tissues Controlling for Cellular Composition and DNAm Batch. (A) Horvath-pan tissue, (B) Hannum, (C) PhenoAge, (D) GrimAge2, (E) DunedinPACE, (F) Skin and Blood, and (G) PedBE (children only). For all clocks, within-person correlations were residualized for tissue cellular composition and DNA methylation batch. * indicate correlations with p<0.05.

## 4. Discussion

We found significant within-person differences in epigenetic clock estimates across most tissues. The largest differences were observed when comparing oral-based versus blood-based tissues (i.e., buccal vs. DBS), however, differences within oral-based (i.e., buccal vs. saliva) and blood-based (i.e., DBS vs. PBMCs) were observed in some clocks. Depending on the clock, buccal and saliva varied significantly in epigenetic age estimates in both magnitude and direction. DBS and buffy coat showed no significant differences in age estimates in all clocks tested. Conversely, DBS and PBMCs displayed significant differences in every clock except the Skin and Blood clock, with DBS having older epigenetic age estimates than PBMCs. As predicted, the Horvath pan-tissue clock showed no significant differences in saliva-DBS and saliva-buffy coat estimates, whereas all other clocks showed significant differences in these comparisons. The Horvath pan-tissue clock did, however, exhibit significant differences between saliva and PBMC, with saliva having higher age estimates. These results were generally consistent when using PC clocks or when controlling for cellular composition and DNAm batch.

Our findings are in line with previous research comparing cross-tissue concordance of epigenetic clocks. A worldwide meta-analysis of epigenetic age acceleration using Horvath pan-tissue, Hannum, PhenoAge and GrimAge2 reported average acceleration values for buccal (27.5 years), saliva (8.6 years), and blood (0.5 years) tissues, indicating comparable acceleration in age estimates for oral-relative to blood-based tissues, as reported here. However, these results should be interpreted with caution because comparisons were made both between-tissue and between-person (Yusipov et al. 2024). A study using pediatric clocks found blood-based tissue correlations ranging between 0.38-0.44, though again, these comparisons were made across different pediatric clocks each calculated in their corresponding tissue (i.e., Knight clock in cord blood and Lee clock in placenta), obscuring inference about tissue-specific effects (Fang et al. 2023). Another study of 21 adults reported within-person correlations between saliva and blood DunedinPoAm (a previous version of DunedinPACE) measurements of 0.60 (all samples) and 0.85 (samples from the same batch) (Raffington et al. 2021). Overall, our results are similar to previous work on the comparability of epigenetic clock estimates across tissues. Here, we extended prior work by providing within-person comparisons of epigenetic clocks across commonly collected tissues in individuals aged 9 to 70 years.

Our findings are also comparable to prior research investigating cross-tissue alignment of alternative genomic measures of biological aging. Our previous work, conducted within this same sample, observed similarly increased biological age, via shorter telomere length (TL), in buccal and saliva relative to blood-based tissues (Wolf et al. 2024). Similar findings were reported in a large-scale meta-analysis, observing stronger correlations between TL among related tissues, e.g., blood-based tissues (McLester-Davis et al. 2023). Previous work has further demonstrated significant differences in quality metrics of DNA across different tissues (Hansen et al. 2007; Lucena-Aguilar et al. 2016; Wolf et al. 2024), however, it remains uncertain to what degree variation in the integrity, purity, and quantity of extracted DNA may influence the reliability of data generated on the EPIC array.

We acknowledge several limitations. First, buffy coat was only collected in children and PBMCs were only collected in adults, limiting comparability of these tissues to selected age ranges. Second, the EPIC v2 DNAm array was used for data collection of all tissue samples. This is the most recent version of the Illumina Infinium arrays and, as it is relatively new, no clocks investigated here were constructed using this array. Though the EPIC v2 array does not contain every probe used in the current epigenetic clock algorithms (see supplementary **Table S1**), recent work has shown the EPIC v1 and EPIC v2 arrays to be highly comparable (Kaur et al. 2023). To increase comparability, imputations for missing probes were made over the entire sample (see *Methods 2.4* and **Supplementary Materials**) (Sugden 2023). Third, tissue samples of children included in the current study were collected from a high-risk pediatric cohort (Schreier et al. 2021) following youth with and without recent investigations for suspected child maltreatment exposure. Although child maltreatment has been shown to alter global and specific gene DNAm levels (Parade et al. 2021), past work with a subset of this cohort has shown that child maltreatment is not associated with changes in epigenetic clock measurements (Etzel et al. 2022), thereby reducing the chance that differences in child maltreatment exposure had an effect on our results. Finally, correlations within-person and between-tissue clock estimates are only informative if DNAm measurements are technically reliable. While previous work on the reliability of DNAm measurements has been reported in blood-based samples (Sugden et al. 2020), the reliability of DNAm measurements of oral-based tissues is less understood. However, one study found high intraclass correlation coefficients (>0.73) between 24 technical replicates of buccal tissue for various epigenetic clocks (Raffington et al. 2023), but further work is needed to determine the reliability of DNAm measurements in these tissues. Therefore, correlations between blood-based and oral-based tissue clock estimates could be limited by the reliability of oral-based DNAm measurements.

Our study suggests that epigenetic clocks can be most reliably applied within the tissue(s) used to generate each clock. Caution should be taken both in research and commercial settings to ensure proper tissue samples are collected for the intended epigenetic clocks. Both research and commercial efforts of measuring biological age using epigenetic clocks may exhibit inaccurate age estimates if incorrect tissues are used. In addition, we recommend the construction and utilization of epigenetic clocks trained on oral-based tissues, thereby enabling the reliable and sensitive estimates of epigenetic age in less invasive tissue types. Overall, our work suggests that tissue type plays an important role in the estimation of biological age and should be carefully considered when using epigenetic clocks.

## Acknowledgements

We thank the children and caregivers for their participation in the study, Child Health Study staff, all nurses at the CRC and the adult participants in this study.

## Data Availability

The datasets used and/or analyzed during the current study are available from the corresponding author upon reasonable request.

## Code Availability

Code used to compute epigenetic clock estimates, “golden standard” datasets, cellular proportions of tissues and perform statistical analyses is available at https://github.com/abnerapsley1/CrossTissueEpiClock

