## Supplemental Materials for "Cross-Tissue Comparison of Epigenetic Aging Clocks in Humans"

### Section S1. Supplementary Methods and Materials

#### S1.1. EPIC v1 and EPIC v2 Illumina Methylation Arrays

Most epigenetic clocks have been constructed using the probes present on either the 27k, 450k, or EPIC v1 Illumina chips. However, due to the recent development and release of the EPIC v2 chip, some probes used to generate epigenetic clocks are either missing or duplicated in the EPIC v2 chip. All samples in our study were assayed using the EPIC v2 array, therefore, comparisons between the two arrays were not made in the current study.

Although we did not use any EPIC v1 arrays in the current project, we present the following information to promote accurate comparability between epigenetic clocks generated using EPIC v1 and EPIC v2 arrays. To compare epigenetic clocks calculated with EPIC v1 and EPIC v2 arrays, two main concerns need to be addressed:

1. How should probes in epigenetic clock algorithms that contain duplicate probes on the EPIC v2 chip be addressed?
2. How should probes in epigenetic clock algorithms that are missing on the EPIC v2 chip be addressed?

##### *S1.1.1 – Duplicated Probes*

In order to remove duplicate probes in the EPIC v2 chip, we used the R package *ENmix* (using v1.36.08 or newer). This package has a function called *rm.cgsuffix*. This function both removes the suffix on EPIC v2 probe names (i.e., removes "_BC11" from "cg25324105_BC11") and collapses duplicate probes by taking their averages.

##### *S1.1.2 – Missing Probes*

When EPIC v2 chips are used, each epigenetic clock has a specific number of probes that are missing from the original algorithm. For example, the DunedinPACE algorithm is missing 29 of its original 173 probes used. Currently, the solution to these missing probes is to impute them. The word imputation is used loosely here, because if a probe is missing in all samples, the process of estimating the probe values is technically considered "extrapolation". Regardless, the values of these missing probes are imputed/extrapolated by being set equal to the mean of a "golden standard" (GS) dataset, with each dataset differing by the clock being used. This means that all samples receive the exact same value (the mean of the GS dataset) for missing probes, thereby decreasing variation in the estimated values of the epigenetic clock.

#### S1.2. "Golden Standard" Datasets for Imputation

In our current work, we have opted to use the imputation/extrapolation method mentioned above, where the probe values are replaced with the mean of the GS dataset probes. However, because there are different GS datasets used for each clock, and each of these datasets is based on a single tissue type, we generated custom GS datasets stratified by tissue and age. This resulted in a total of 8 GS datasets: adult buccal, children buccal, adult saliva, children saliva, adult whole blood, children whole blood, adult PBMCs and children PBMCs. Each GS dataset was constructed using 2 publicly available datasets, one using the EPIC v1 array and another using the 450k array. All datasets with available IDAT files were preprocessed using the same procedures described in the manuscript (see *Methods*). By necessity, all datasets with raw intensity files available excluded the bisulfite conversion and bead count filtering steps. All publicly available datasets were BMIQ normalized before the generation of GS datasets. (See **Table S2** for dataset details).

### S1.3. Estimations of Cellular Composition

Because cell types have unique DNA methylation signatures, the cellular composition of a tissue sample can be a confounding factor when estimating epigenetic clocks. We performed estimations of cellular composition for each tissue in the current study using two distinct methods.

#### *S1.3.1 – The Houseman-Blood Method*

The first method, which we refer to as the "Houseman-Blood" method was used for all blood-based tissues (dry blood spots (DBS), buffy coat, and peripheral blood mononuclear cells (PBMCs)). The Houseman-Blood method, which has been widely used for estimating immune cell compositions and has been detailed previously (Houseman et al. 2012), estimates cellular proportions of CD8T, CD4T, natural killer cells, B-cells, monocytes and granulocytes from DNA methylation data. These estimates are then used when controlling for cellular compositions of blood-based tissues.

#### *S1.3.2 – The Houseman-ReferenceFree Method*

The second method, which we refer to as the "Houseman-ReferenceFree" method was used for buccal and saliva tissue. The Houseman-ReferenceFree method can be used for any type of tissue (including non-blood tissue) and will provide estimates for cellular proportions based on DNA methylation data and a user-defined input that specifies the number of cell types to estimate (Houseman et al. 2016). In addition to providing cellular proportions, the Houseman-ReferenceFree method also provides a protocol for estimating the "best-fitting" number of cell subtypes in a provided sample. Briefly, deviance statistics are computed for each possible number of cell types present (2 cell types, 3 cell types, … ). These deviance statistics are computed for a random selection of the study sample a defined number of times (we used 1000 bootstrapped samples). Then, summary statistics for each possible number of cell types present are compared and the number of cell types with the minimum deviance summary statistic are chosen.

We performed these calculations with our buccal and saliva samples separately. We extracted the 10,000 probes with the highest variation in buccal and saliva separately and subsequently used these probes in cellular composition estimates. After computation, we determined that buccal tissue contained 3 cellular subtypes and saliva contained 5 (see **Figure S1**).

**Section S2. Supplementary Results and Figures**

S2.1. Supplementary Tables

| Epigenetic Clock | Total CpG Probes | Number of Probes Present in EPIC v2 | Number of Probes Absent in EPIC v2 | Proportion Missing |
|---|---|---|---|---|
| Horvath | 353 | 340 | 13 | 3.68% |
| Hannum | 71 | 64 | 7 | 9.86% |
| PhenoAge | 513 | 495 | 18 | 3.51% |
| GrimAge2 | 1030 | 845 | 185 | 17.96% |
| DunedinPACE | 173 | 144 | 29 | 16.76% |
| PedBE | 94 | 93 | 1 | 1.06% |
| Skin and Blood | 391 | 374 | 17 | 4.35% |

Table S1: An overview of the number of CpG probes included in the EPIC v2 array for each clock used in the present analysis.

| GEO Accession Number | Tissue | Sample Number | Sample Inclusion Criteria | Age | Array | Raw IDAT Used | Intensity Matrix Used | Sample Pval Filter | Sample Bisulfite Converstion Filter | Probe Bead Count Filter | Probe Pval Filter | Normalization |
|---|---|---|---|---|---|---|---|---|---|---|---|---|
| GSE50586 | Buccal | 10 | Only controls included | Adult | 450k | No | Yes | Yes | No | No | Yes | BMIQ |
| GSE166844 | Buccal | 28 | Only buccal samples | Adult | EPIC v1 | No | Yes | Yes | No | No | Yes | BMIQ |
| GSE50759 | Buccal | 40 | Only controls included | Children | 450k | No | Yes | Yes | No | No | Yes | BMIQ |
| GSE147058 | Buccal | 48 | Only twin A included | Children | EPIC v1 | Yes | No | Yes | Yes | Yes | Yes | BMIQ |
| GSE245924 | PBMC | 16 | Only controls included | Adult | EPIC v1 | Yes | No | Yes | Yes | Yes | Yes | BMIQ |
| GSE117929 | PBMC | 19 | Only controls included | Adult | 450k | No | Yes | Yes | No | No | Yes | BMIQ |
| GSE132181 | PBMC | 50 | Only PBMC age 7 included, used first 50 samples | Children | EPIC v1 | Yes | No | Yes | Yes | Yes | Yes | BMIQ |
| GSE40576 | PBMC | 30 | Only first 30 controls included | Children | 450k | No | Yes | Yes | No | No | Yes | BMIQ |
| GSE130153 | Saliva | 22 | Only saliva test samples included | Adult | 450k | No | Yes | Yes | No | No | Yes | BMIQ |
| GSE111165 | Saliva | 15 | Only EPIC saliva samples over 18 | Adult | EPIC v1 | Yes | No | Yes | Yes | Yes | Yes | BMIQ |
| GSE147318 | Saliva | 18 | Only whole saliva samples | Children | EPIC v1 | Yes | No | Yes | Yes | Yes | Yes | BMIQ |
| GSE112314 | Saliva | 22 | All samples included | Children | 450k | Yes | No | Yes | Yes | Yes | Yes | BMIQ |
| GSE218186 | Whole Blood | 55 | Only controls included | Adult | EPIC v1 | Yes | No | Yes | Yes | Yes | Yes | BMIQ |
| GSE201287 | Whole Blood | 40 | Only controls included | Adult | 450k | Yes | No | Yes | Yes | Yes | Yes | BMIQ |
| GSE221864 | Whole Blood | 31 | Only controls included | Children | EPIC v1 | Yes | No | Yes | Yes | Yes | Yes | BMIQ |
| GSE174555 | Whole Blood | 17 | Only controls included | Children | 450k | Yes | No | Yes | Yes | Yes | Yes | BMIQ |

Table S2: A description of the public datasets and preprocessing methods used to construct custom “golden standard” (GS) datasets for imputation of missing CpGs.

| | Buccal | | | Saliva | | | DBS | | | Buffy Coat | | | PBMC | | |
|---|---|---|---|---|---|---|---|---|---|---|---|---|---|---|---|
| | **Mean** | **Median** | **SD** | **Mean** | **Median** | **SD** | **Mean** | **Median** | **SD** | **Mean** | **Median** | **SD** | **Mean** | **Median** | **SD** |
| **Horvath** | 32.08 | 24.84 | 20.67 | 43.44 | 36.43 | 21.59 | 41.13 | 34.56 | 20.12 | 23.05 | 22.52 | 4.48 | 53.74 | 53.23 | 16.87 |
| **Horvath Acceleration** | 1.60 | 1.13 | 4.39 | 11.15 | 11.28 | 7.38 | 13.12 | 13.17 | 4.07 | 11.18 | 11.02 | 3.87 | 9.99 | 10.26 | 4.95 |
| **Hannum** | 41.76 | 39.03 | 15.18 | 41.16 | 42.60 | 15.46 | 14.46 | 8.82 | 18.36 | -1.73 | -1.17 | 3.88 | 24.27 | 23.24 | 15.12 |
| **Hannum Acceleration** | 11.27 | 12.03 | 7.77 | 8.87 | 9.53 | 12.88 | -13.56 | -13.58 | 4.52 | -13.6 | -13.25 | 3.44 | -19.47 | -19.54 | 5.57 |

Table S3: The mean, median and standard deviation for the Horvath pan-tissue and Hannum clock raw estimates and acceleration estimates.

| | Horvath Pan-Tissue Acceleration | | | | | | | Hannum Acceleration | | | | | | |
|---|---|---|---|---|---|---|---|---|---|---|---|---|---|---|
| | **ICC** | **F** | **df1** | **df2** | **P-value** | **Lower Bound** | **Upper Bound** | **ICC** | **F** | **df1** | **df2** | **P-value** | **Lower Bound** | **Upper Bound** |
| **ICC1** | 0.09 | 1.5 | 82 | 332 | 0.009 | 0.01 | 0.18 | -0.05 | 0.76 | 82 | 332 | 0.930 | -0.10 | 0.02 |
| **ICC2** | 0.17 | 3.1 | 82 | 328 | <0.001 | 0.06 | 0.30 | 0.11 | 5.29 | 82 | 328 | <0.001 | 0.01 | 0.25 |
| **ICC3** | 0.30 | 3.1 | 82 | 328 | <0.001 | 0.20 | 0.41 | 0.46 | 5.29 | 82 | 328 | <0.001 | 0.36 | 0.57 |
| **ICC1k** | 0.32 | 1.5 | 82 | 332 | 0.009 | 0.06 | 0.53 | -0.31 | 0.76 | 82 | 332 | 0.930 | -0.82 | 0.09 |
| **ICC2k** | 0.50 | 3.1 | 82 | 328 | <0.001 | 0.23 | 0.68 | 0.38 | 5.29 | 82 | 328 | <0.001 | 0.06 | 0.62 |
| **ICC3k** | 0.68 | 3.1 | 82 | 328 | <0.001 | 0.56 | 0.78 | 0.81 | 5.29 | 82 | 328 | <0.001 | 0.74 | 0.87 |

Table S4: Intraclass correlation coefficients across all tissues for the Horvath-Pan Tissue and Hannum acceleration estimates. ICC1 - One-way random effects, absolute agreement, single rater/measurement, ICC2 - Two-way random effects, absolute agreement, single rater/measurement, ICC3 - Two-way mixed effects, consistency, single rater/measurement, ICC1k - One-way random effects, absolute agreement, multiple raters/measurements, ICC2k - Two-way random effects, absolute agreement, multiple raters/measurements, ICC3k - Two-way mixed effects, consistency, multiple raters/measurements. F – Fisher's F statistic, df – degrees of freedom.

| | Buccal | | | Saliva | | | DBS | | | Buffy Coat | | | PBMC | | |
|---|---|---|---|---|---|---|---|---|---|---|---|---|---|---|---|
| | **Mean** | **Median** | **SD** | **Mean** | **Median** | **SD** | **Mean** | **Median** | **SD** | **Mean** | **Median** | **SD** | **Mean** | **Median** | **SD** |
| **PhenoAge** | 43.94 | 36.16 | 18.85 | 50.31 | 49.60 | 22.98 | 21.7 | 15.01 | 22.17 | 3.42 | 3.93 | 5.83 | 29.03 | 26.21 | 18.91 |
| **PhenoAge Acceleration** | 13.45 | 14.13 | 6.19 | 18.02 | 18.04 | 12.27 | -6.31 | -6.87 | 5.98 | -8.44 | -7.36 | 5.38 | -14.72 | -14.14 | 6.68 |
| **GrimAge2** | 71.68 | 67.68 | 15.36 | 59.17 | 57.63 | 15.27 | 34.69 | 30.37 | 14.50 | 23.14 | 23.28 | 3.89 | 42.01 | 41.66 | 12.73 |
| **GrimAge2 Acceleration** | 41.19 | 42.88 | 8.10 | 26.88 | 25.86 | 13.36 | 6.67 | 8.23 | 5.73 | 11.27 | 11.15 | 3.43 | -1.74 | -1.99 | 6.15 |

Table S5: The mean, median and standard deviation for the PhenoAge and GrimAge2 clock raw estimates and acceleration estimates.

| | PhenoAge Acceleration | | | | | | | GrimAge2 Acceleration | | | | | | |
|---|---|---|---|---|---|---|---|---|---|---|---|---|---|---|
| | **ICC** | **F** | **df1** | **df2** | **P-value** | **Lower Bound** | **Upper Bound** | **ICC** | **F** | **df1** | **df2** | **P-value** | **Lower Bound** | **Upper Bound** |
| **ICC1** | -0.13 | 0.45 | 82 | 332 | 1.000 | -0.16 | -0.08 | -0.05 | 0.76 | 82 | 332 | 0.930 | -0.10 | 0.02 |
| **ICC2** | 0.05 | 2.39 | 82 | 328 | <0.001 | 0.00 | 0.12 | 0.12 | 7.61 | 82 | 328 | <0.001 | 0.01 | 0.27 |
| **ICC3** | 0.22 | 2.39 | 82 | 328 | <0.001 | 0.13 | 0.33 | 0.57 | 7.61 | 82 | 328 | <0.001 | 0.47 | 0.67 |
| **ICC1k** | -1.25 | 0.45 | 82 | 332 | 1.000 | -2.11 | -0.56 | -0.31 | 0.76 | 82 | 332 | 0.930 | -0.82 | 0.08 |
| **ICC2k** | 0.21 | 2.39 | 82 | 328 | <0.001 | 0.01 | 0.40 | 0.40 | 7.61 | 82 | 328 | <0.001 | 0.06 | 0.64 |
| **ICC3k** | 0.58 | 2.39 | 82 | 328 | <0.001 | 0.42 | 0.71 | 0.87 | 7.61 | 82 | 328 | <0.001 | 0.82 | 0.91 |

Table S6: Intraclass correlation coefficients across all tissues for the PhenoAge and GrimAge2 acceleration estimates. ICC1 - One-way random effects, absolute agreement, single rater/measurement, ICC2 - Two-way random effects, absolute agreement, single rater/measurement, ICC3 - Two-way mixed effects, consistency, single rater/measurement, ICC1k - One-way random effects, absolute agreement, multiple raters/measurements, ICC2k - Two-way random effects, absolute agreement, multiple raters/measurements, ICC3k - Two-way mixed effects, consistency, multiple raters/measurements. F – Fisher's F statistic, df – degrees of freedom.

| | Buccal | | | Saliva | | | DBS | | | Buffy Coat | | | PBMC | | |
|---|---|---|---|---|---|---|---|---|---|---|---|---|---|---|---|
| | **Mean** | **Median** | **SD** | **Mean** | **Median** | **SD** | **Mean** | **Median** | **SD** | **Mean** | **Median** | **SD** | **Mean** | **Median** | **SD** |
| **PACE** | 1.61 | 1.62 | 0.08 | 1.50 | 1.52 | 0.25 | 0.96 | 0.96 | 0.09 | 0.96 | 0.96 | 0.09 | 0.89 | 0.89 | 0.10 |

Table S7: The mean, median and standard deviation for the DunedinPACE estimates.

| | DunedinPACE | | | | | | |
|---|---|---|---|---|---|---|---|
| | **ICC** | **F** | **df1** | **df2** | **P-value** | **Lower Bound** | **Upper Bound** |
| **ICC1** | -0.17 | 0.27 | 82 | 332 | 1.000 | -0.19 | -0.14 |
| **ICC2** | 0.03 | 2.28 | 82 | 328 | <0.001 | 0.00 | 0.08 |
| **ICC3** | 0.20 | 2.28 | 82 | 328 | <0.001 | 0.11 | 0.31 |
| **ICC1k** | -2.74 | 0.27 | 82 | 332 | 1.000 | -4.19 | -1.61 |
| **ICC2k** | 0.13 | 2.28 | 82 | 328 | <0.001 | -0.01 | 0.29 |
| **ICC3k** | 0.56 | 2.28 | 82 | 328 | <0.001 | 0.39 | 0.70 |

Table S8: Intraclass correlation coefficients across all tissues for the DunedinPACE estimates. ICC1 - One-way random effects, absolute agreement, single rater/measurement, ICC2 - Two-way random effects, absolute agreement, single rater/measurement, ICC3 - Two-way mixed effects, consistency, single rater/measurement, ICC1k - One-way random effects, absolute agreement, multiple raters/measurements, ICC2k - Two-way random effects, absolute agreement, multiple raters/measurements, ICC3k - Two-way mixed effects, consistency, multiple raters/measurements. F – Fisher's F statistic, df – degrees of freedom.

| | Buccal | | | Saliva | | | DBS | | | Buffy Coat | | | PBMC | | |
|---|---|---|---|---|---|---|---|---|---|---|---|---|---|---|---|
| | **Mean** | **Median** | **SD** | **Mean** | **Median** | **SD** | **Mean** | **Median** | **SD** | **Mean** | **Median** | **SD** | **Mean** | **Median** | **SD** |
| **Skin and Blood** | 28.65 | 21.29 | 20.32 | 32.09 | 26.81 | 20.80 | 25.39 | 15.91 | 19.45 | 8.95 | 8.86 | 1.44 | 40.48 | 41.98 | 16.25 |
| **Skin and Blood Acceleration** | -1.84 | -1.91 | 3.41 | -0.21 | -0.35 | 6.20 | -2.63 | -2.77 | 2.36 | -2.92 | -3.09 | 1.25 | -3.26 | -3.23 | 3.53 |
| **PedBE** | 12.89 | 13.11 | 1.77 | 9.32 | 9.31 | 1.69 | 7.07 | 7.12 | 0.55 | 7.03 | 6.92 | 0.49 | -- | -- | -- |
| **PedBE Acceleration** | 0.97 | 1.06 | 1.25 | -2.86 | -2.83 | 1.63 | -4.97 | -5.01 | 1.12 | -4.84 | -4.74 | 1.14 | -- | -- | -- |

Table S9: The mean, median and standard deviation for the Skin and Blood and PedBE clock raw estimates and acceleration estimates.

| | Skin and Blood Acceleration | | | | | | | PedBE Acceleration | | | | | | |
|---|---|---|---|---|---|---|---|---|---|---|---|---|---|---|
| | **ICC** | **F** | **df1** | **df2** | **P-value** | **Lower Bound** | **Upper Bound** | **ICC** | **F** | **df1** | **df2** | **P-value** | **Lower Bound** | **Upper Bound** |
| **ICC1** | 0.18 | 2.1 | 82 | 332 | <0.001 | 0.09 | 0.28 | 0.67 | 11 | 82 | 332 | <0.001 | 0.58 | 0.75 |
| **ICC2** | 0.19 | 2.3 | 82 | 328 | <0.001 | 0.11 | 0.30 | 0.68 | 42 | 82 | 328 | <0.001 | 0.34 | 0.84 |
| **ICC3** | 0.21 | 2.3 | 82 | 328 | <0.001 | 0.12 | 0.32 | 0.89 | 42 | 82 | 328 | <0.001 | 0.85 | 0.92 |
| **ICC1k** | 0.52 | 2.1 | 82 | 332 | <0.001 | 0.33 | 0.66 | 0.91 | 11 | 82 | 332 | <0.001 | 0.87 | 0.94 |
| **ICC2k** | 0.54 | 2.3 | 82 | 328 | <0.001 | 0.37 | 0.68 | 0.92 | 42 | 82 | 328 | <0.001 | 0.72 | 0.96 |
| **ICC3k** | 0.57 | 2.3 | 82 | 328 | <0.001 | 0.40 | 0.70 | 0.98 | 42 | 82 | 328 | <0.001 | 0.97 | 0.98 |

Table S10: Intraclass correlation coefficients across all tissues for the Skin and Blood and PedBE acceleration estimates. ICC1 - One-way random effects, absolute agreement, single rater/measurement, ICC2 - Two-way random effects, absolute agreement, single rater/measurement, ICC3 - Two-way mixed effects, consistency, single rater/measurement, ICC1k - One-way random effects, absolute agreement, multiple raters/measurements, ICC2k - Two-way random effects, absolute agreement, multiple raters/measurements, ICC3k - Two-way mixed effects, consistency, multiple raters/measurements. F – Fisher's F statistic, df – degrees of freedom.

| | Buccal vs Saliva | | Buccal vs DBS | | Buccal vs PBMC | | Saliva vs DBS | | Saliva vs PBMC | | DBS vs PBMC | |
|---|---|---|---|---|---|---|---|---|---|---|---|---|
| | **Estimate** | **p-value** | **Estimate** | **p-value** | **Estimate** | **p-value** | **Estimate** | **p-value** | **Estimate** | **p-value** | **Estimate** | **p-value** |
| **Horvath** | -9.79 | <0.001 | -11.76 | <0.001 | -8.12 | <0.001 | -2.41 | 0.160 | 2.22 | 0.183 | 4.33 | <0.001 |
| **Hannum** | 2.81 | 0.017 | 23.89 | <0.001 | 28.26 | <0.001 | 20.63 | <0.001 | 24.16 | <0.001 | 5.46 | <0.001 |
| **PhenoAge** | -6.48 | 0.002 | 17.99 | <0.001 | 27.10 | <0.001 | 23.79 | <0.001 | 32.75 | <0.001 | 11.23 | <0.001 |
| **GrimAge2** | 16.43 | <0.001 | 35.89 | <0.001 | 40.25 | <0.001 | 17.45 | <0.001 | 22.42 | <0.001 | 5.16 | <0.001 |
| **DunedinPACE** | 0.15 | <0.001 | 0.66 | <0.001 | 0.75 | <0.001 | 0.47 | <0.001 | 0.58 | <0.001 | 0.1 | <0.001 |
| **Skin and Blood** | -2.59 | 0.041 | 1.52 | 0.027 | 2.01 | 0.005 | 3.66 | 0.031 | 3.98 | 0.008 | 0.86 | 0.115 |

Table S11: Paired t-test results for within-person, between-tissue *adult* clock comparisons. This table mirrors **Table 2** from the manuscript. Positive values indicate a larger clock estimate for the tissue listed first in the tissue pair (i.e., a positive buccal vs. saliva value indicates buccal having a higher estimate than saliva).

| | Buccal vs Saliva | | Buccal vs DBS | | Buccal vs Buffy Coat | | Saliva vs DBS | | Saliva vs Buffy Coat | | DBS vs Buffy Coat | |
|---|---|---|---|---|---|---|---|---|---|---|---|---|
| | **Estimate** | **p-value** | **Estimate** | **p-value** | **Estimate** | **p-value** | **Estimate** | **p-value** | **Estimate** | **p-value** | **Estimate** | **p-value** |
| **Horvath** | -8.86 | <0.001 | -10.28 | <0.001 | -10.12 | <0.001 | -1.06 | 0.361 | -1.69 | 0.196 | -0.27 | 0.614 |
| **Hannum** | -1.31 | 0.600 | 28.96 | <0.001 | 29.29 | <0.001 | 27.40 | <0.001 | 29.28 | <0.001 | 0.02 | 0.978 |
| **PhenoAge** | -3.35 | 0.160 | 24.56 | <0.001 | 23.98 | <0.001 | 25.71 | <0.001 | 24.93 | <0.001 | -1.33 | 0.085 |
| **GrimAge2** | 8.39 | <0.001 | 35.66 | <0.001 | 34.82 | <0.001 | 25.82 | <0.001 | 25.70 | <0.001 | -1.11 | 0.181 |
| **DunedinPACE** | 0.01 | 0.905 | 0.63 | <0.001 | 0.62 | <0.001 | 0.62 | <0.001 | 0.61 | <0.001 | -0.02 | 0.217 |
| **Skin and Blood** | -0.76 | 0.255 | 1.10 | <0.001 | 0.84 | 0.009 | 1.58 | 0.036 | 1.32 | 0.055 | -0.29 | 0.123 |

Table S12: Paired t-test results for within-person, between-tissue children clock comparisons. This table mirrors **Table 2** from the manuscript. Positive values indicate a larger clock estimate for the tissue listed first in the tissue pair (i.e., a positive buccal vs. saliva value indicates buccal having a higher estimate than saliva).

| | Buccal vs Saliva | | Buccal vs DBS | | Buccal vs Buffy Coat | | Buccal vs PBMC | | Saliva vs DBS | | Saliva vs Buffy Coat | | Saliva vs PBMC | | DBS vs Buffy Coat | | DBS vs PBMC | |
|---|---|---|---|---|---|---|---|---|---|---|---|---|---|---|---|---|---|---|
| | **Est.** | **Pvalue** | **Est.** | **Pvalue** | **Est.** | **Pvalue** | **Est.** | **Pvalue** | **Est.** | **Pvalue** | **Est.** | **Pvalue** | **Est.** | **Pvalue** | **Est.** | **Pvalue** | **Est.** | **Pvalue** |
| **HorvathPC** | -11.41 | <0.001 | -18.90 | <0.001 | -13.56 | <0.001 | -18.21 | <0.001 | -8.12 | <0.001 | -8.54 | <0.001 | -2.38 | 0.110 | -0.25 | 0.420 | 6.00 | <0.001 |
| **HannumPC** | 7.54 | <0.001 | 28.70 | <0.001 | 32.55 | <0.001 | 34.18 | <0.001 | 19.65 | <0.001 | 25.46 | <0.001 | 24.73 | <0.001 | -0.11 | 0.800 | 9.54 | <0.001 |
| **PhenoAgePC** | 14.31 | <0.001 | 59.60 | <0.001 | 64.13 | <0.001 | 68.60 | <0.001 | 42.51 | <0.001 | 50.03 | <0.001 | 50.68 | <0.001 | -0.64 | 0.360 | 14.41 | <0.001 |
| **GrimAgePC** | 7.75 | <0.001 | 20.01 | <0.001 | 20.58 | <0.001 | 25.24 | <0.001 | 11.42 | <0.001 | 13.53 | <0.001 | 16.22 | <0.001 | -0.61 | 0.038 | 6.67 | <0.001 |
| **Skin and Blood PC** | -6.60 | <0.001 | -10.34 | <0.001 | -10.56 | <0.001 | -1.51 | 0.190 | -4.07 | <0.001 | -5.27 | <0.001 | 5.76 | <0.001 | -0.49 | 0.290 | 9.33 | <0.001 |

Table S13: Paired t-test results for within-person, between-tissue PC clock comparisons. This table mirrors **Table 2** from the manuscript. Positive values indicate a larger clock estimate for the tissue listed first in the tissue pair (i.e., a positive buccal vs. saliva value indicates buccal having a higher estimate than saliva).

| | Buccal | | Saliva | | DBS | | Buffy Coat | | PBMC | |
|---|---|---|---|---|---|---|---|---|---|---|
| | **Estimate** | **p-value** | **Estimate** | **p-value** | **Estimate** | **p-value** | **Estimate** | **p-value** | **Estimate** | **p-value** |
| **HorvathPC** | 13.11 | <0.001 | 11.63 | <0.001 | 4.70 | <0.001 | 2.59 | <0.001 | 8.04 | <0.001 |
| **HannumPC** | -27.14 | <0.001 | -20.94 | <0.001 | -24.54 | <0.001 | -24.57 | <0.001 | -20.55 | <0.001 |
| **PhenoAgePC** | -40.11 | <0.001 | -19.91 | <0.001 | -1.74 | 0.005 | -0.94 | 0.160 | 1.71 | 0.042 |
| **GrimAgePC** | 7.39 | <0.001 | 1.52 | 0.084 | -8.00 | <0.001 | -6.97 | <0.001 | -6.88 | <0.001 |
| **Skin and Blood PC** | -3.45 | <0.001 | -8.34 | <0.001 | -15.37 | <0.001 | -17.01 | <0.001 | -5.25 | <0.001 |

Table S14: Paired t-test results comparing the standard and PC clock estimates within-person and within-tissue. Positive values indicate a larger standard clock estimate and negative values indicate a larger PC clock estimate.

S2.2. Estimation of Tissue Cellular Composition

**Figure S1** shows the results from using the Houseman-ReferenceFree method on both buccal and saliva tissue to estimate the "best-fitting" number of cell subtypes. 1000 bootstrapped samples composed of around 36.8% of the original sample were drawn randomly from the tissue of interest (with replacement) and deviance statistics were calculated for each sample using an assumed 1-15 cell subtypes. Code for the calculation of deviance statistics is located at https://github.com/abnerapsley1/CrossTissueEpiClock.

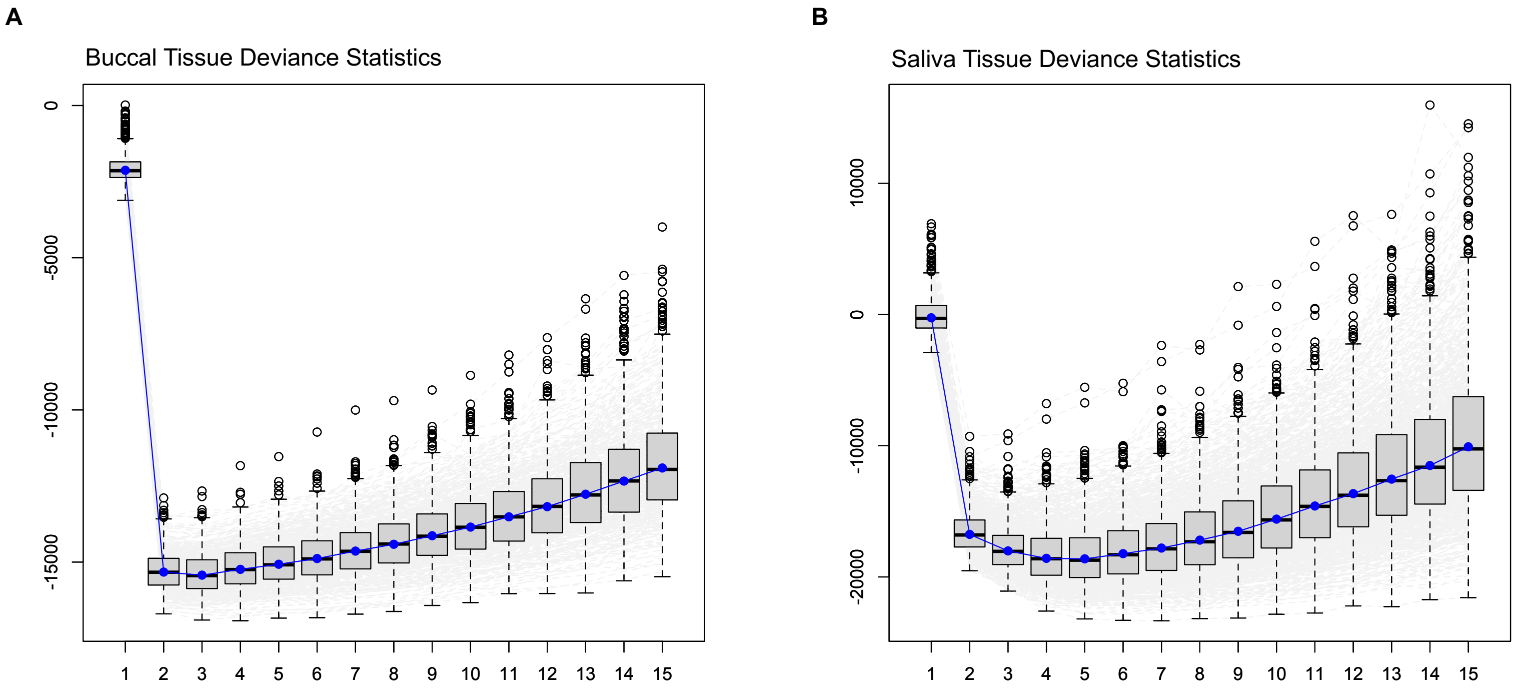


Figure S1 – Houseman-ReferenceFree Method Buccal (A) and Saliva (B) Deviance Statistics. X-axes show the number of estimated cell types and y-axes show the deviance statistic. Blue dots indicate the median deviance statistic, grey boxes indicate upper and lower quartiles and whiskers indicate 95$^{th}$ percentiles. An estimation of three cell types best fit the buccal tissue (A) and five cell types best fit the saliva tissue (B).

S2.3. Comparability of First-Generation Epigenetic Clocks Across Tissues – Stratified by Age

**Figure S2** parallels **Figure 1** from the main text, however only adults (18 years or older) are included in the sample. Similarly, **Figure S3** parallels **Figure 1** from the main text but only includes children (younger than 18 years old).

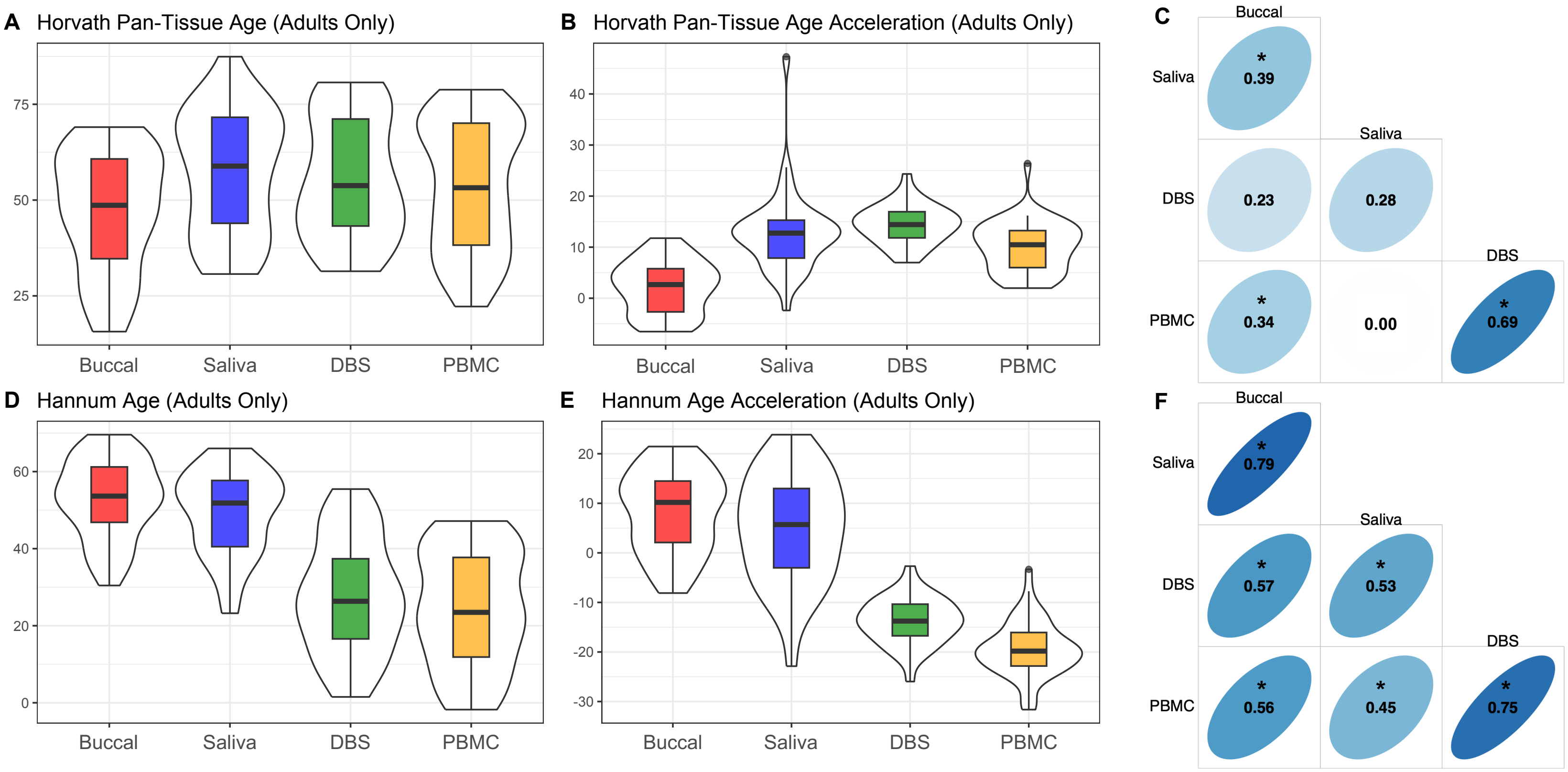


Figure S2 – First Generation Epigenetic Clock Distributions and Correlations (Adults Only).
(A) Horvath pan-tissue epigenetic age estimates, (B) Horvath pan-tissue epigenetic age acceleration estimates, (C) Within-person Horvath pan-tissue epigenetic age acceleration correlations across tissues, (D) Hannum epigenetic age estimates, (E) Hannum

epigenetic age acceleration estimates and (F) Within-person Hannum epigenetic age acceleration correlations across tissues. Thick black horizontal bars on violin plots indicate the tissue-stratified median clock value and colored boxes indicate interquartile ranges.

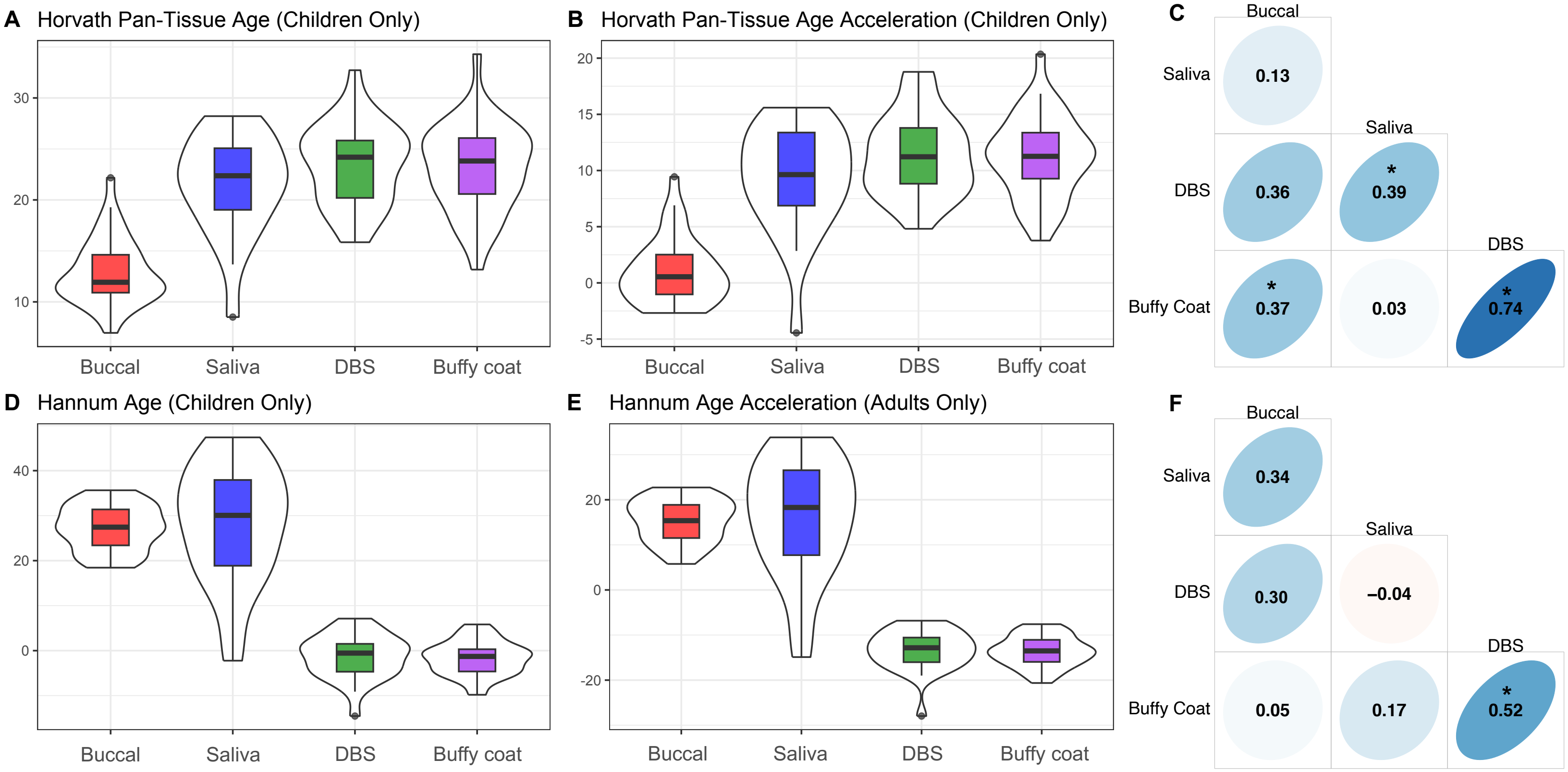


Figure S3 – First Generation Epigenetic Clock Distributions and Correlations (Children Only).
(A) Horvath pan-tissue epigenetic age estimates, (B) Horvath pan-tissue epigenetic age acceleration estimates, (C) Within-person Horvath pan-tissue epigenetic age acceleration correlations across tissues, (D) Hannum epigenetic age estimates, (E) Hannum epigenetic age acceleration estimates and (F) Within-person Hannum epigenetic age acceleration correlations across tissues. Thick black horizontal bars on violin plots indicate the tissue-stratified median clock value and colored boxes indicate interquartile ranges.

S2.4. Comparability of Second-Generation Epigenetic Clocks Across Tissues – Stratified by Age

**Figure S4** parallels **Figure 2** from the main text, however only adults (18 years or older) are included in the sample. Similarly, **Figure S5** parallels **Figure 2** from the main text but only includes children (younger than 18 years old).

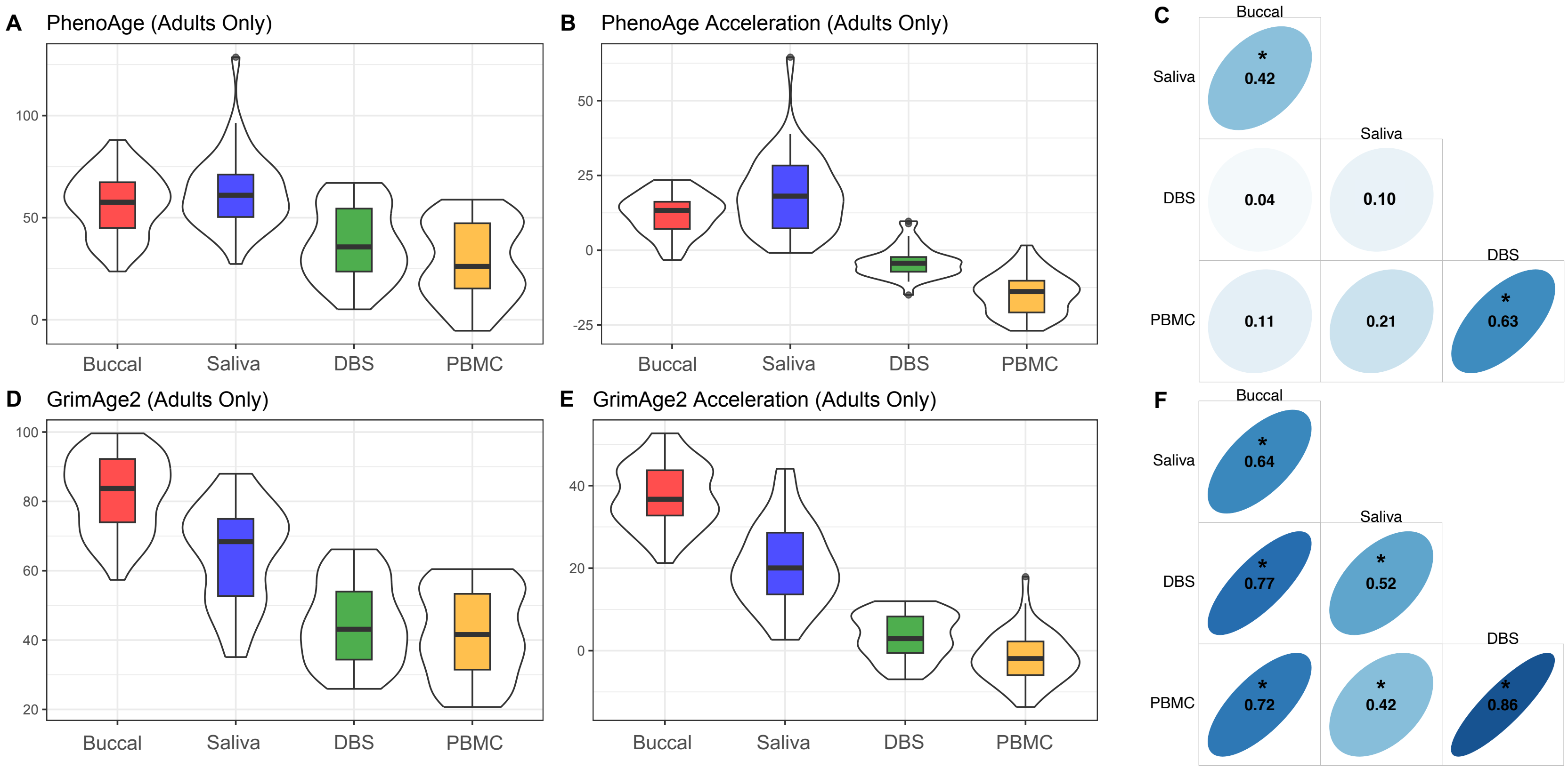


Figure S4 – Second Generation Epigenetic Clock Distributions and Correlations (Adults Only). (A) PhenoAge epigenetic age estimates, (B) PhenoAge epigenetic age acceleration estimates, (C) Within-person PhenoAge epigenetic age acceleration correlations across tissues, (D) GrimAge2 epigenetic age estimates, (E) GrimAge2 epigenetic age acceleration estimates and (F) Within-person GrimAge2 epigenetic age acceleration correlations across tissues. Thick black horizontal bars on violin plots indicate the tissue-stratified median clock value and colored boxes indicate interquartile ranges.

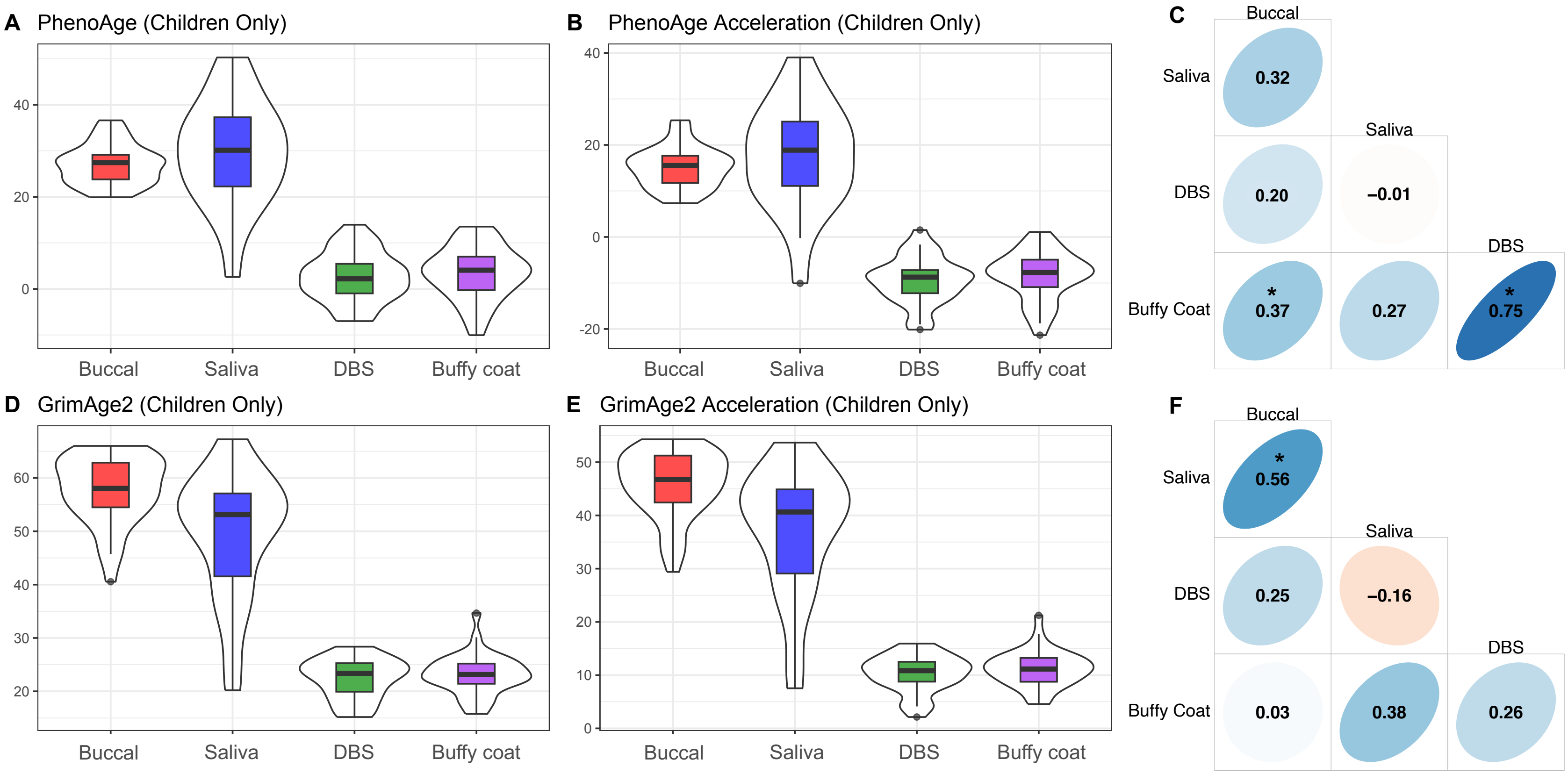


Figure S5 - Second Generation Epigenetic Clock Distributions and Correlations (Children Only).
(A) PhenoAge epigenetic age estimates, (B) PhenoAge epigenetic age acceleration estimates, (C) Within-person PhenoAge epigenetic age acceleration correlations across tissues, (D) GrimAge2 epigenetic age estimates, (E) GrimAge2 epigenetic age acceleration estimates and (F) Within-person GrimAge2 epigenetic age acceleration correlations across tissues. Thick black horizontal bars on violin plots indicate the tissue-stratified median clock value and colored boxes indicate interquartile ranges.

S2.5. Comparability of DunedinPACE Epigenetic Clock Across Tissues – Stratified by Age

**Figure S6** parallels **Figure 3** from the main text, however only adults (18 years or older) are included in the sample. Similarly, **Figure S7** parallels **Figure 3** from the main text but only includes children (younger than 18 years old).

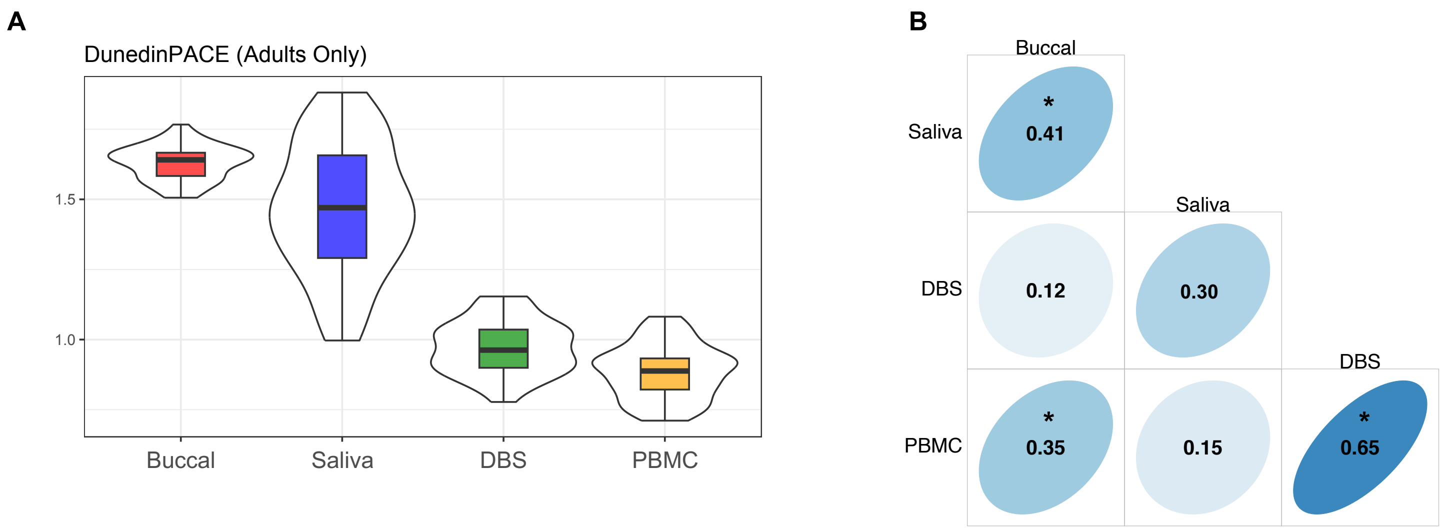


Figure S6 - DunedinPace Epigenetic Clock Distributions and Correlations (Adults Only).
(A) DunedinPACE estimates, (B) Within-person DunedinPACE estimate correlations across tissues. Thick black horizontal bars on violin plots indicate the tissue-stratified median clock value and colored boxes indicate interquartile ranges.

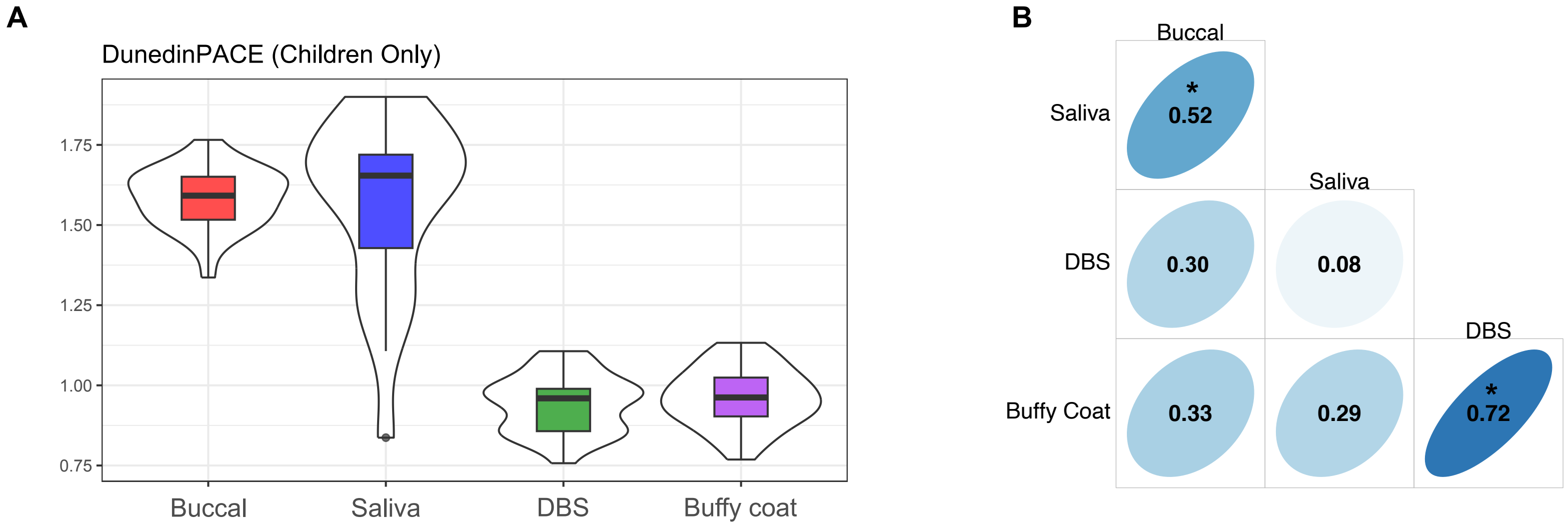


Figure S7 - DunedinPace Epigenetic Clock Distributions and Correlations (Children Only).
(A) DunedinPACE estimates, (B) Within-person DunedinPACE estimate correlations across tissues. Thick black horizontal bars on violin plots indicate the tissue-stratified median clock value and colored boxes indicate interquartile ranges.

S2.6. Comparability of Skin/Blood and PedBE Epigenetic Clocks Across Tissues – Stratified by Age

**Figure S8** parallels **Figure 4** from the main text, however only adults (18 years or older) are included in the sample and the PedBE clock is not included. Similarly, **Figure S9** parallels **Figure 4** from the main text but only includes children (younger than 18 years old) and does not include the PedBE clock.

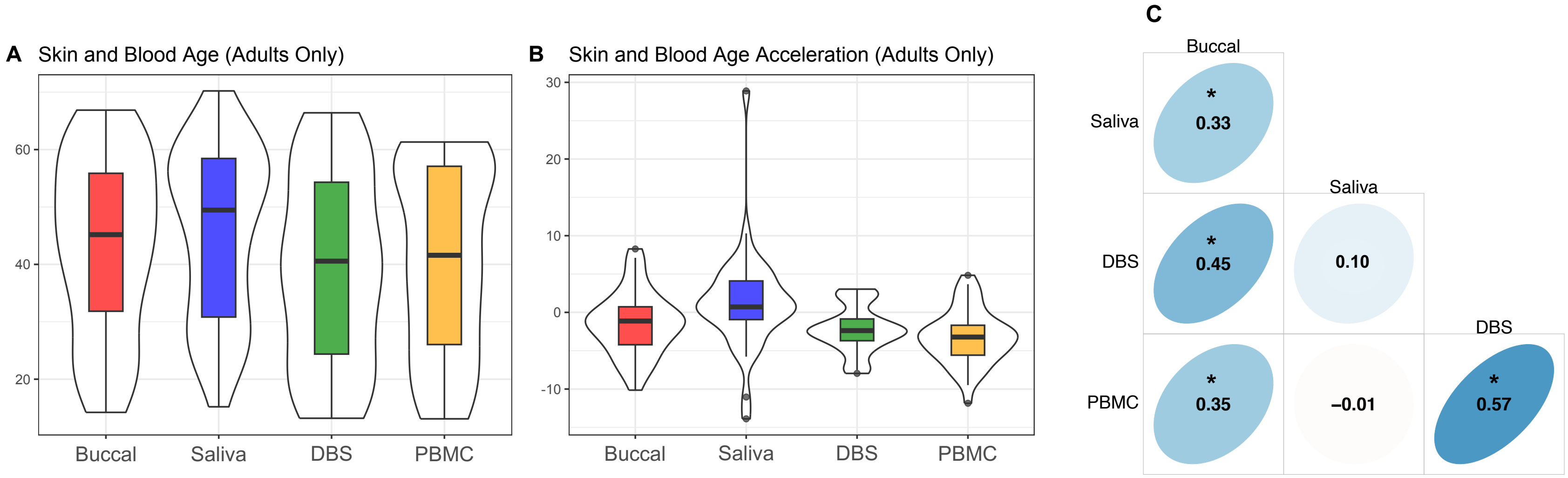


Figure S8 – Skin/Blood Clock Distributions and Correlations (Adults Only).
(A) Skin/Blood epigenetic age estimates, (B) Skin/Blood epigenetic age acceleration estimates, (C) Within-person Skin/Blood epigenetic age acceleration correlations across tissues. Thick black horizontal bars on violin plots indicate the tissue-stratified median clock value and colored boxes indicate interquartile ranges.

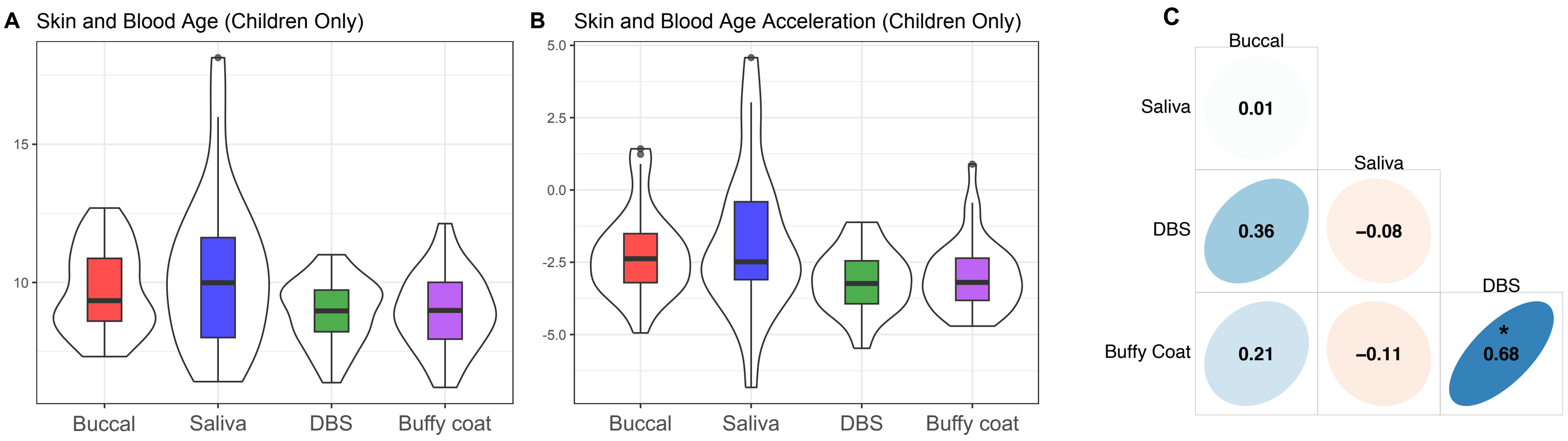


Figure S9 – Skin/Blood Clock Distributions and Correlations (Children Only).
(A) Skin/Blood epigenetic age estimates, (B) Skin/Blood epigenetic age acceleration estimates, (C) Within-person Skin/Blood epigenetic age acceleration correlations across tissues. Thick black horizontal bars on violin plots indicate the tissue-stratified median clock value and colored boxes indicate interquartile ranges.

S2.7. Sensitivity Analyses – PC Clocks (Full Sample and Stratified by Age)
The following figures are full sample (**Figures S10-S12**) and age-stratified results (**Figures S13-S18**) of principal component clocks.

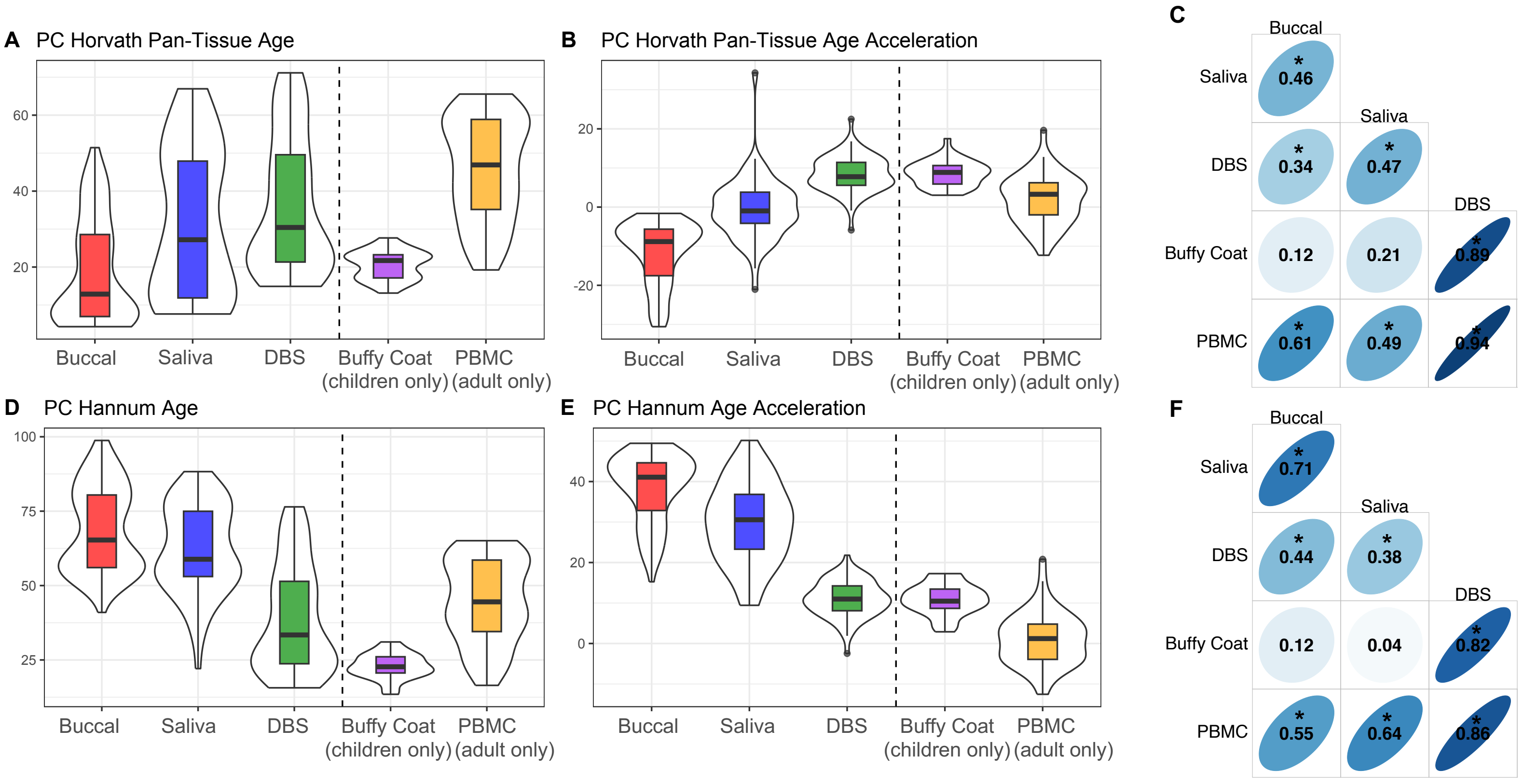


Figure S10 – First Generation Principal Component Epigenetic Clock Distributions and Correlations.
(A) PC-Horvath pan-tissue epigenetic age estimates, (B) PC-Horvath pan-tissue epigenetic age acceleration estimates, (C) Within-person PC-Horvath pan-tissue epigenetic age acceleration correlations across tissues, (D) PC-Hannum epigenetic age estimates, (E) PC-Hannum epigenetic age acceleration estimates and (F) Within-person PC-Hannum epigenetic age acceleration correlations across

tissues. Thick black horizontal bars on violin plots indicate the tissue-stratified median clock value and colored boxes indicate interquartile ranges.

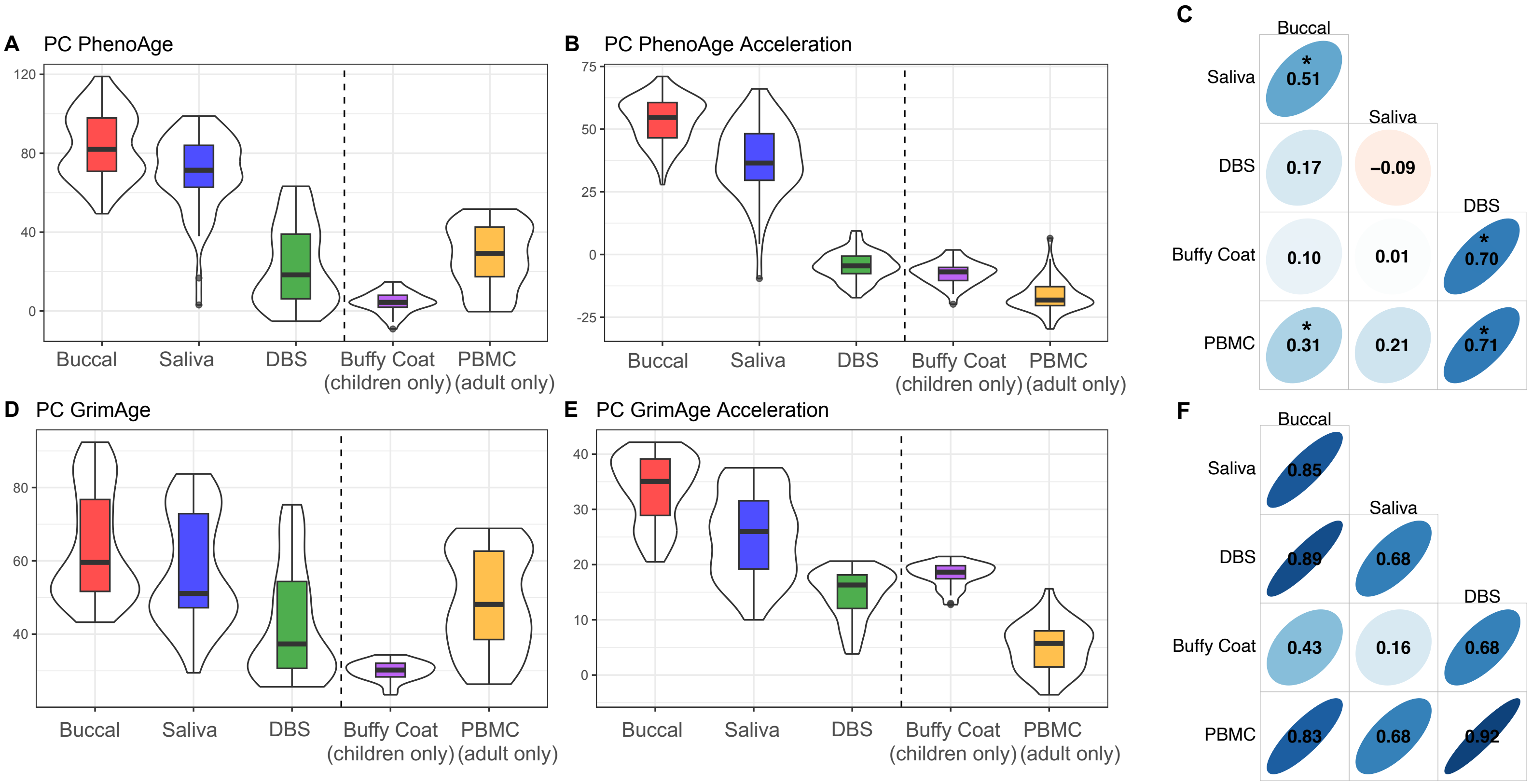


Figure S11 – Second Generation Principal Component Epigenetic Clock Distributions and Correlations. (A) PC-PhenoAge epigenetic age estimates, (B) PC-PhenoAge epigenetic age acceleration estimates, (C) Within-person PC-PhenoAge epigenetic age acceleration correlations across tissues, (D) PC-GrimAge epigenetic age estimates, (E) PC-GrimAge epigenetic age acceleration estimates and (F) Within-person PC-GrimAge epigenetic age acceleration correlations across tissues. Thick black horizontal bars on violin plots indicate the tissue-stratified median clock value and colored boxes indicate interquartile ranges.

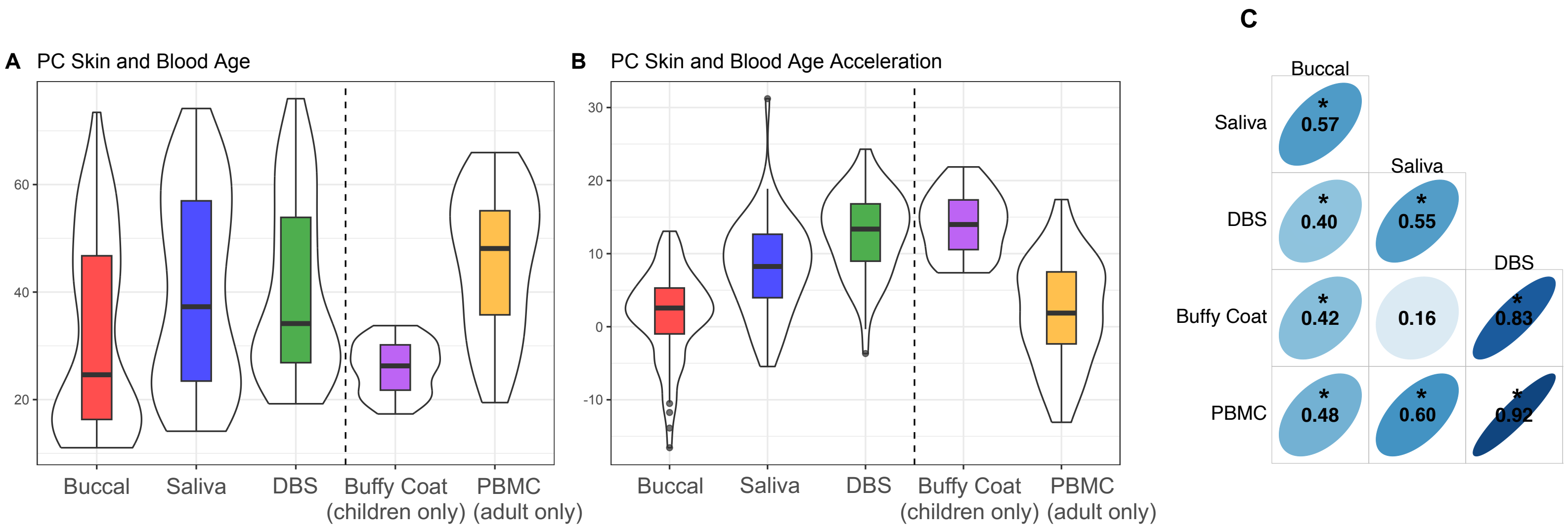


Figure S12 – Skin and Blood Principal Component Epigenetic Clock Distributions and Correlations.
(A) PC-SkinBlood epigenetic age estimates, (B) PC-SkinBlood epigenetic age acceleration estimates, (C) Within-person PC-SkinBlood epigenetic age acceleration correlations across tissues. Thick black horizontal bars on violin plots indicate the tissue-stratified median clock value and colored boxes indicate interquartile ranges.

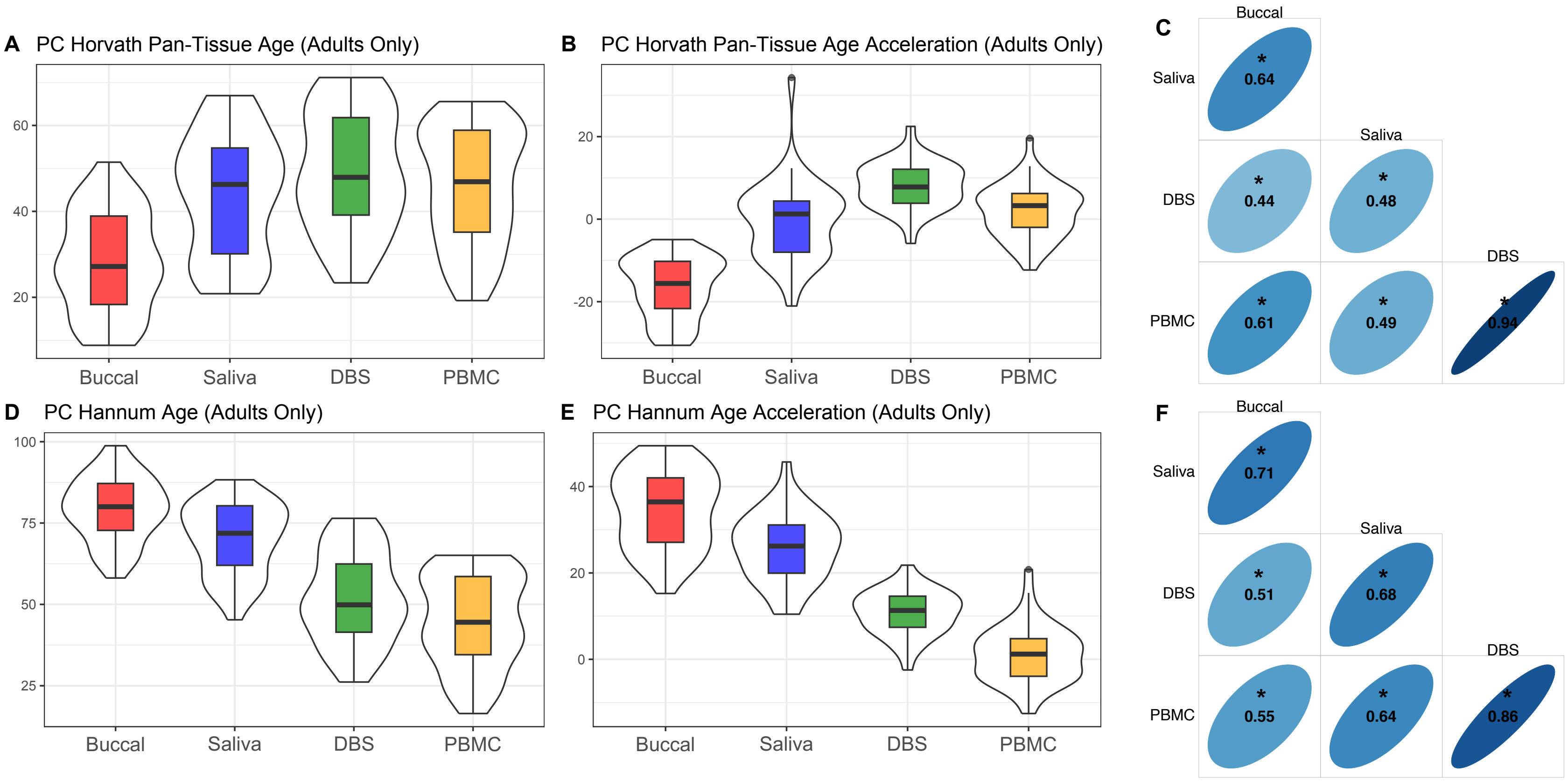


Figure S13 – First Generation Principal Component Epigenetic Clock Distributions and Correlations (Adults Only).
(A) PC-Horvath pan-tissue epigenetic age estimates, (B) PC-Horvath pan-tissue epigenetic age acceleration estimates, (C) Within-person PC-Horvath pan-tissue epigenetic age acceleration correlations across tissues, (D) PC-Hannum epigenetic age estimates, (E) PC-Hannum epigenetic age acceleration estimates and (F) Within-person PC-Hannum epigenetic age acceleration correlations across tissues. Thick black horizontal bars on violin plots indicate the tissue-stratified median clock value and colored boxes indicate interquartile ranges.

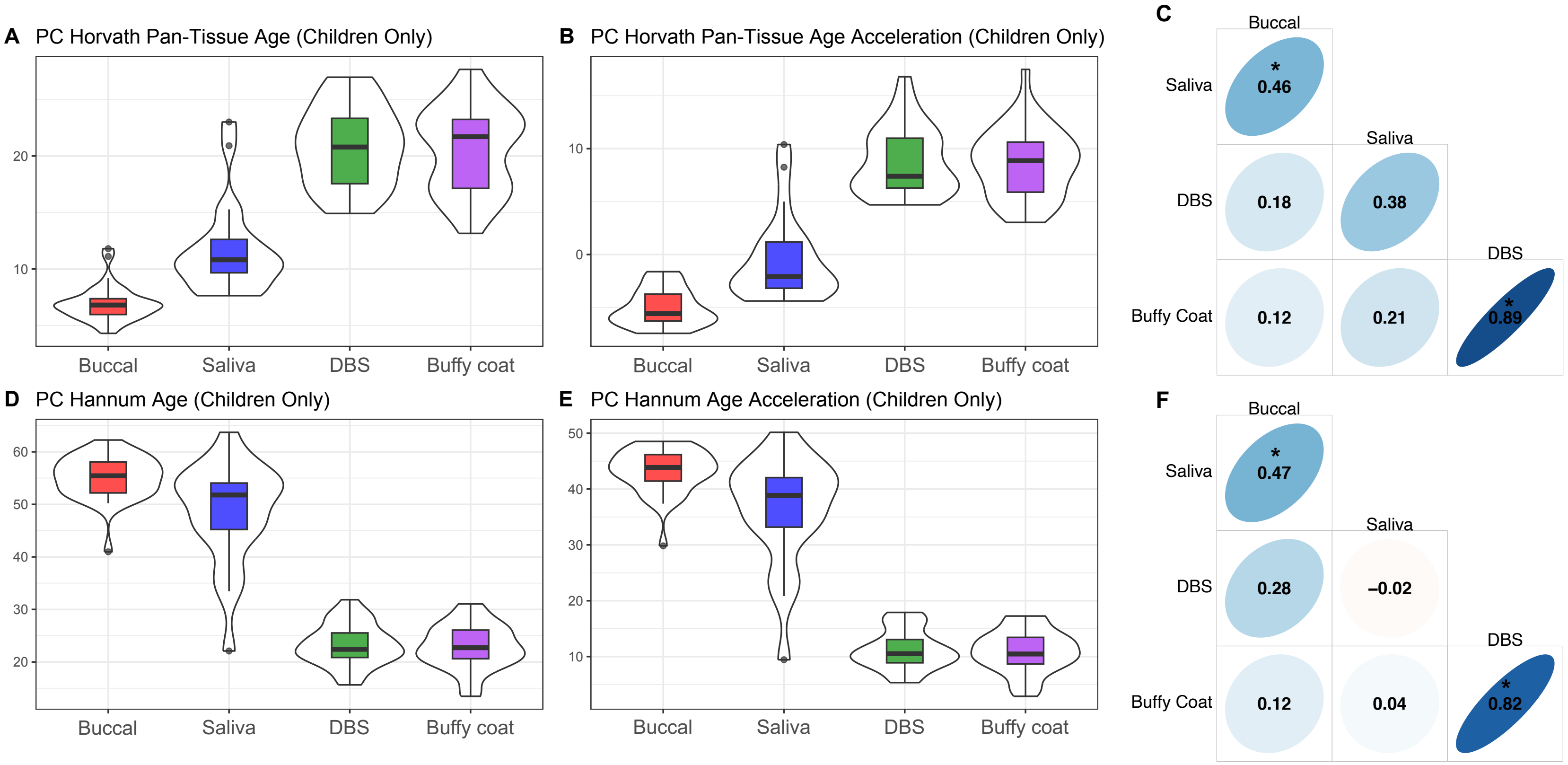


Figure S14 – First Generation Principal Component Epigenetic Clock Distributions and Correlations (Children Only). (A) PC-Horvath pan-tissue epigenetic age estimates, (B) PC-Horvath pan-tissue epigenetic age acceleration estimates, (C) Within-person PC-Horvath pan-tissue epigenetic age acceleration correlations across tissues, (D) PC-Hannum epigenetic age estimates, (E) PC-Hannum epigenetic age acceleration estimates and (F) Within-person PC-Hannum epigenetic age acceleration correlations across tissues. Thick black horizontal bars on violin plots indicate the tissue-stratified median clock value and colored boxes indicate interquartile ranges.

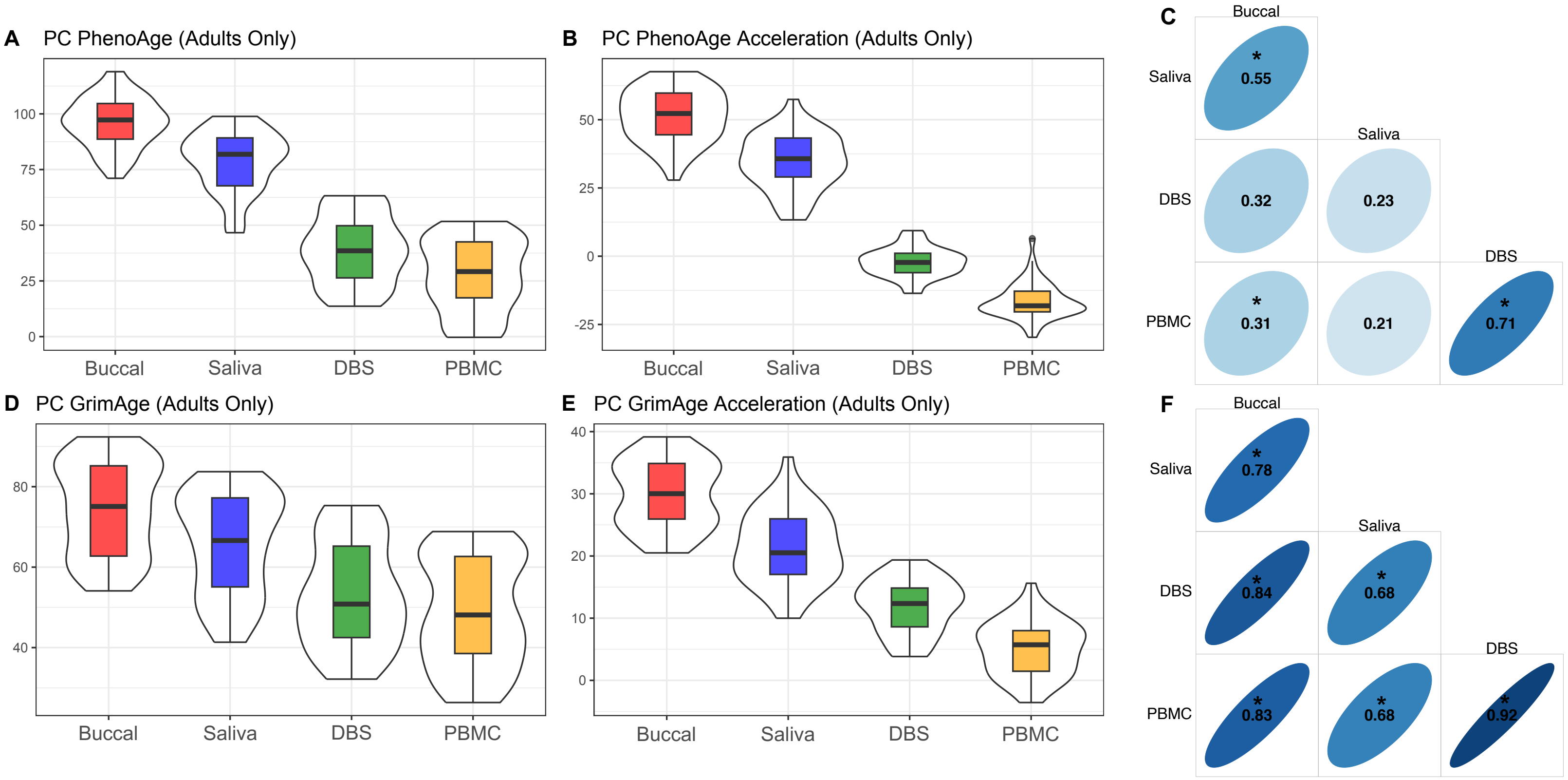


Figure S15 – Second Generation Principal Component Epigenetic Clock Distributions and Correlations (Adults Only). (A) PC-PhenoAge epigenetic age estimates, (B) PC-PhenoAge epigenetic age acceleration estimates, (C) Within-person PC-PhenoAge epigenetic age acceleration correlations across tissues, (D) PC-GrimAge epigenetic age estimates, (E) PC-GrimAge epigenetic age acceleration estimates and (F) Within-person PC-GrimAge epigenetic age acceleration correlations across tissues. Thick black horizontal bars on violin plots indicate the tissue-stratified median clock value and colored boxes indicate interquartile ranges.

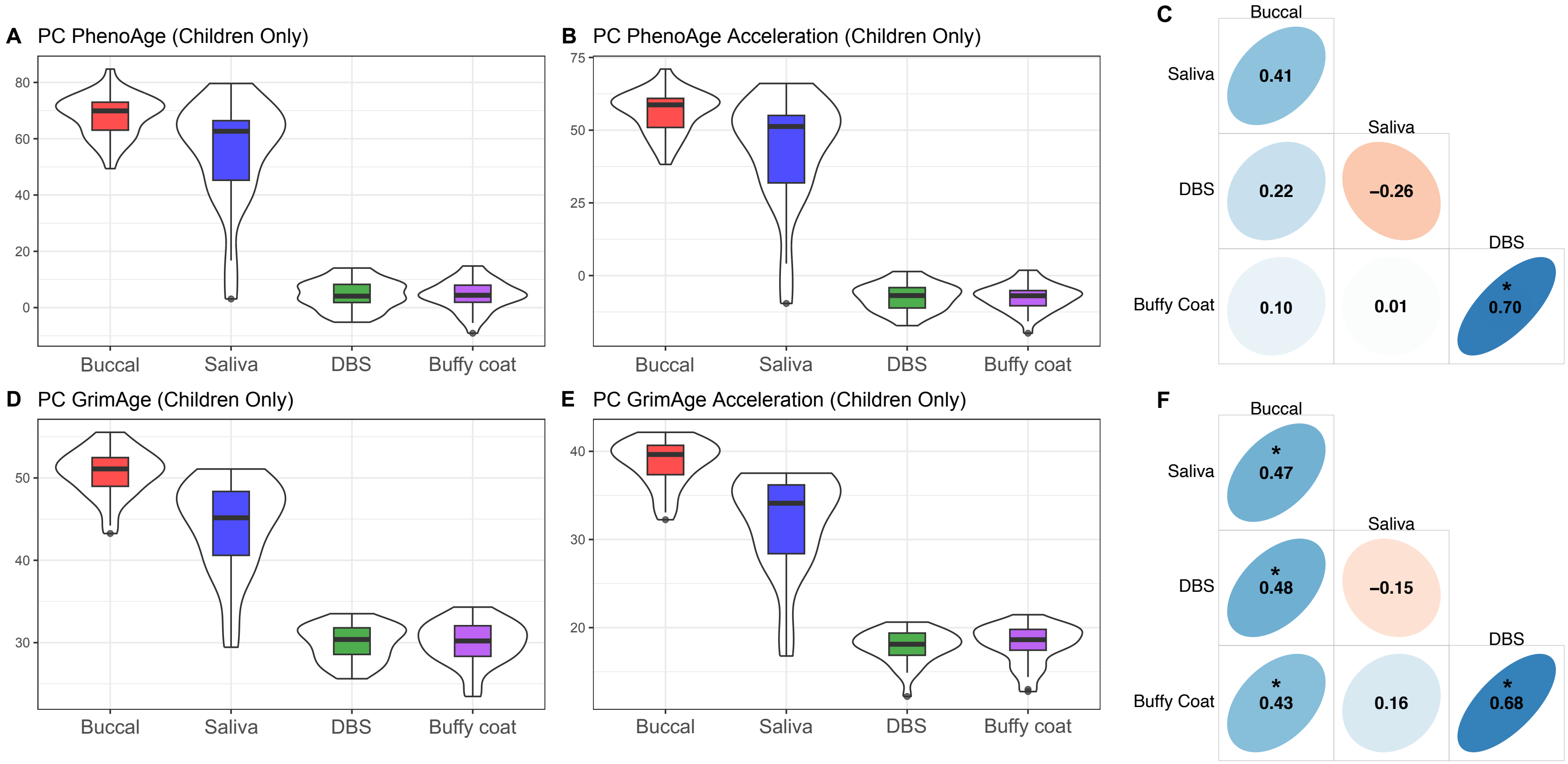


Figure S16 – Second Generation Principal Component Epigenetic Clock Distributions and Correlations (Children Only). (A) PC-PhenoAge epigenetic age estimates, (B) PC-PhenoAge epigenetic age acceleration estimates, (C) Within-person PC-PhenoAge epigenetic age acceleration correlations across tissues, (D) PC-GrimAge epigenetic age estimates, (E) PC-GrimAge epigenetic age acceleration estimates and (F) Within-person PC-GrimAge epigenetic age acceleration correlations across tissues. Thick black horizontal bars on violin plots indicate the tissue-stratified median clock value and colored boxes indicate interquartile ranges.

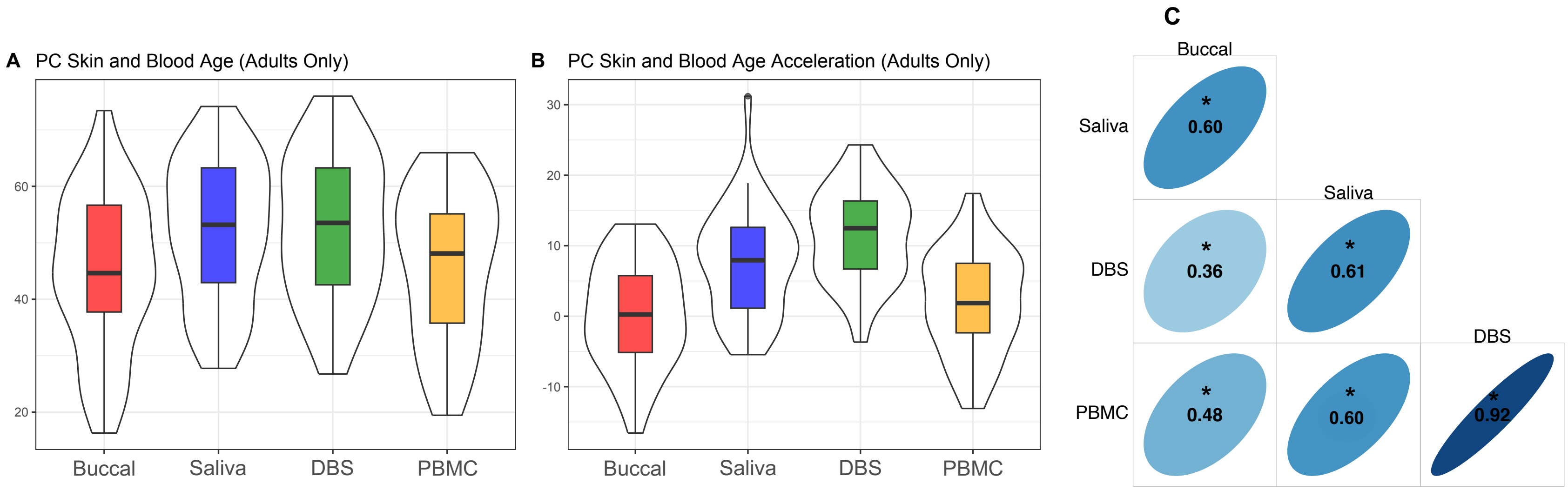


Figure S17 – Skin and Blood Principal Component Epigenetic Clock Distributions and Correlations (Adults Only). (A) PC-SkinBlood epigenetic age estimates, (B) PC-SkinBlood epigenetic age acceleration estimates, (C) Within-person PC-SkinBlood epigenetic age acceleration correlations across tissues. Thick black horizontal bars on violin plots indicate the tissue-stratified median clock value and colored boxes indicate interquartile ranges.

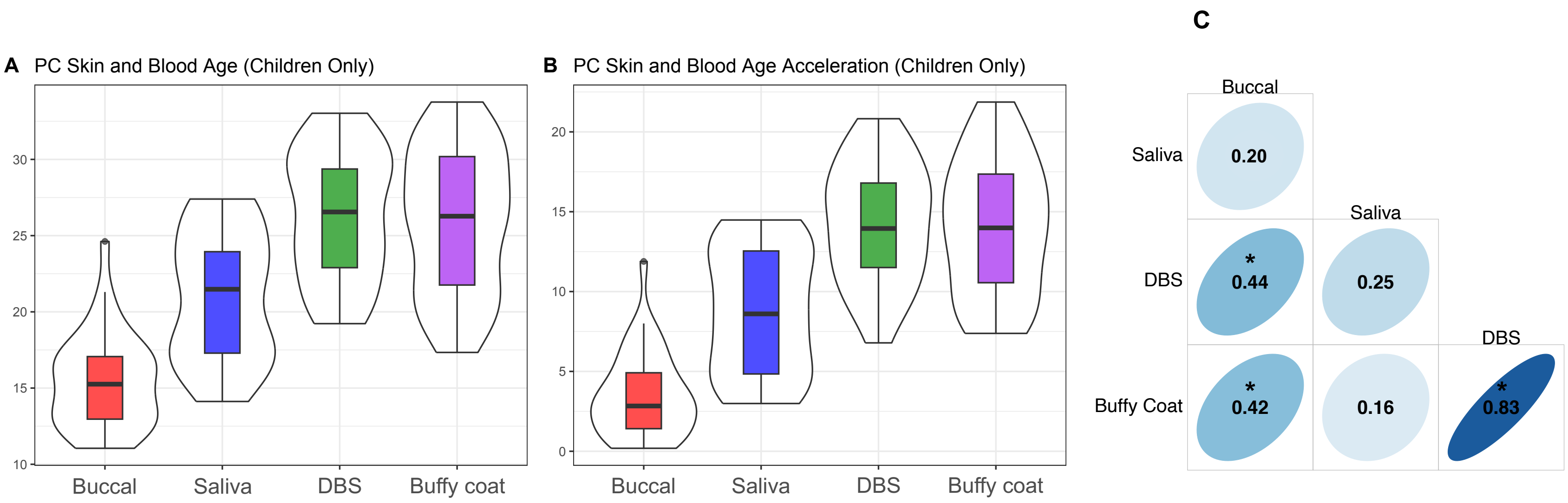


Figure S18 – Skin and Blood Principal Component Epigenetic Clock Distributions and Correlations (Children Only). (A) PC-SkinBlood epigenetic age estimates, (B) PC-SkinBlood epigenetic age acceleration estimates, (C) Within-person PC-SkinBlood epigenetic age acceleration correlations across tissues. Thick black horizontal bars on violin plots indicate the tissue-stratified median clock value and colored boxes indicate interquartile ranges.
